# Columnar processing of border ownership in primate visual cortex

**DOI:** 10.1101/2021.08.06.455427

**Authors:** Tom P. Franken, John H. Reynolds

## Abstract

To understand a visual scene, the brain segregates figures from background by assigning borders to foreground objects. Neurons in primate visual cortex encode which object owns a border (border ownership), but the underlying circuitry is not understood. Here we used multielectrode probes to record from border ownership selective units in different layers in macaque visual area V4 to study the laminar organization and timing of border ownership selectivity.

We find that border ownership selectivity occurs first in deep layer units, in contrast to spike latency for small stimuli in the classical receptive field. Units on the same penetration typically share the preferred side of border ownership, also across layers, similar to orientation preference. Units are often border ownership selective for a range of border orientations, where the preferred sides of border ownership are systematically organized in visual space.

Together our data reveal a columnar organization of border ownership in V4 where the earliest border ownership signals are not simply inherited from upstream areas, but computed by neurons in deep layers, and may thus be part of signals fed back to upstream cortical areas or the oculomotor system early after stimulus onset. The finding that preferred border ownership is clustered and can cover a wide range of spatially contiguous locations, suggests that the asymmetric context integrated by these neurons is provided in a systematically clustered manner, possibly through corticocortical feedback and horizontal connections.

## Introduction

One of the deepest and most enduring mysteries of visual perception is how the brain constructs an internal model of the ever-changing world that falls before our eyes. An essential step in this dynamic process of constructive perception is the assignment of border ownership. Consider panel 1 in Figure 1A. The dashed circle indicates the classical receptive field (cRF) of a hypothetical neuron, straddling the edge of an object. The portion of the edge falling inside the circle is perceived to be owned by the light grey square on the bottom left of the edge. Now consider panel 2 in Figure 1A. The local edge within the circle is identical to that in the left panel but this local edge is now perceived as owned by the dark grey square to the upper right of the edge. This phenomenon is called border ownership. It was first recognized by the Gestalt psychologists in the early 20th century and is beautifully illustrated in Rubin’s famous face-vase illusion (Koffka, 1935; Rubin, 1921). Border ownership represents a fundamental computation in visual perception that is thought to be critical to visual scene segmentation and object recognition (Nakayama et al., 1995). von der Heydt and colleagues discovered neurons in primate visual cortex that are selective for border ownership, most prominently in extrastriate visual areas V2 and V4 (Zhou et al., 2000).

**Figure 1.**
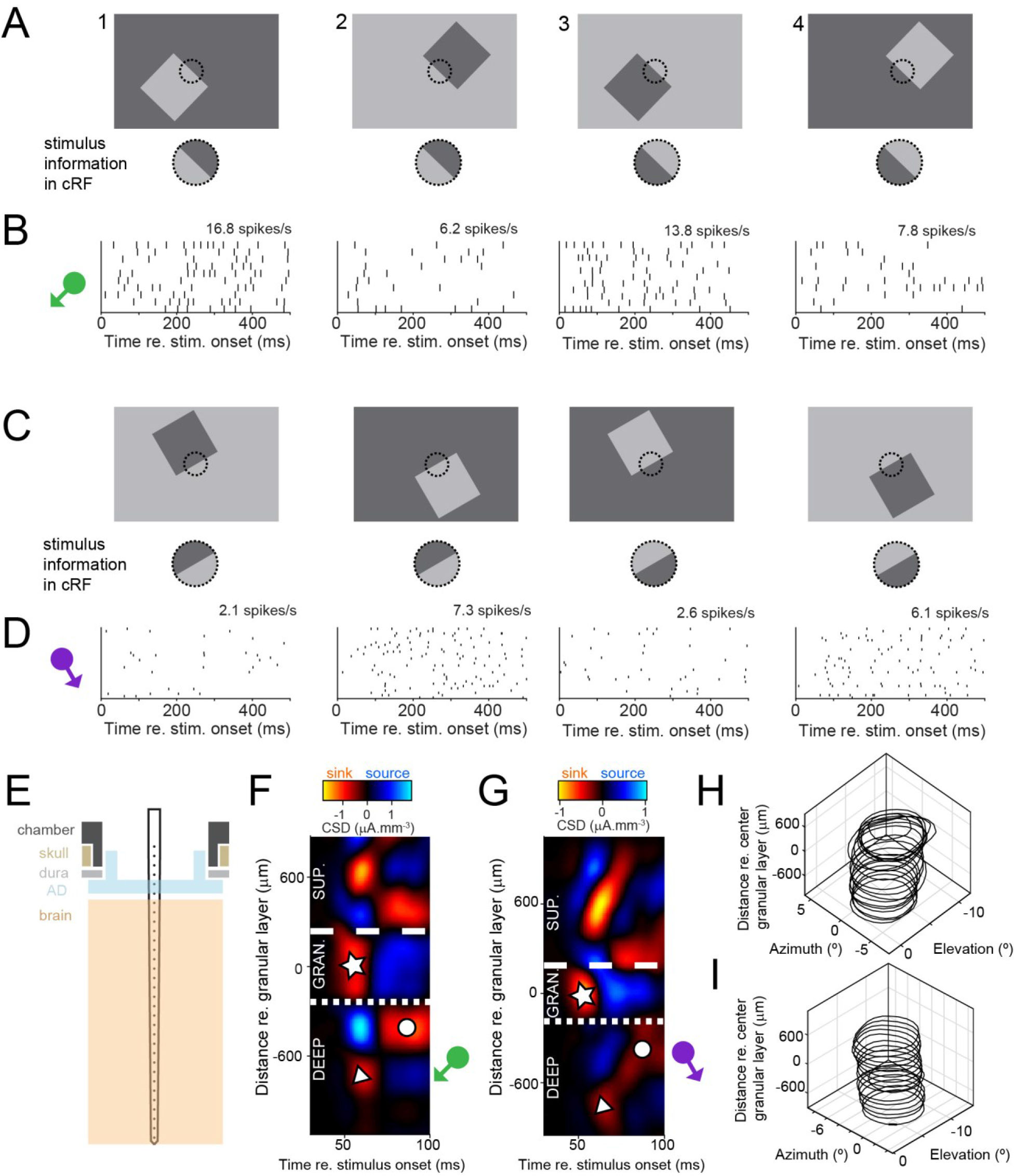
Laminar multielectrode recordings from border ownership selective units in area V4. (**A**) Top row shows set of border ownership stimuli. Black dotted outline represents the classical receptive field (cRF). Bottom row shows that the stimulus information in the cRF is identical for stimuli 1 and 2, and for stimuli 3 and 4. (**B**) Dot rasters showing responses to the stimuli in A from a border-ownership selective well-isolated unit. The symbol on the left indicates the preferred side of border ownership for the unit. Average spike rates during the stimulus window are indicated above the panels. (**C,D**) Similar to A,B, for a multiunit cluster recorded during a different penetration. (**E**) Cartoon showing the recording setup. A laminar multielectrode probe with 32 channels was lowered through a transparent artificial dura (AD), orthogonal relative to the cortical surface. (**F**) Laminar compartments (superficial; granular (input); deep layers) were estimated using current-source density (CSD) analysis. See text for definitions of the compartments and explanation of symbols. Data is from the same penetration during which the unit in A was recorded. The position of the green symbol indicates that the unit shown in B was positioned in the deep layers. See also Figure 1-figure supplement 1 and Figure 1-figure supplement 2. (**G**). Similar to F, for the penetration during which the unit shown in D was recorded. (**H**) Receptive field contours for multiunit activity recorded on different electrode contacts from the penetration shown in F. Contours are drawn at *z* = 3. (**I**) Similar to H, for the penetration shown in G.

Though the existence of border ownership selective neurons has been well established in prior studies using single electrodes (Hesse and Tsao, 2016; O’Herron and von der Heydt, 2009; Zhang and von der Heydt, 2010), how this selectivity arises from cortical circuits remains unclear (Grossberg, 2016; von der Heydt, 2015; Wagatsuma and Sakai, 2017; Yazdanbakhsh and Livingstone, 2006). Some authors have proposed a dominant role for feedforward inputs, which carry information from upstream areas (Sakai et al., 2012; Sakai and Nishimura, 2006; Supèr et al., 2010), whereas others posit a central role for horizontal connections and/or cortical feedback (Craft et al., 2007; Grossberg, 2016; Hu and Niebur, 2017; Zhang and von der Heydt, 2010; Zhaoping, 2005). These pathways have a distinct laminar organization: feedforward inputs arrive primarily in the granular (input) layer, whereas horizontal and cortico-cortical feedback connections predominantly target superficial and deep layers (Douglas and Martin, 2004; Rockland and Pandya, 1979; Rockland et al., 1994; Rockland and Lund, 1983; Ungerleider and Desimone, 1986; Yoshioka et al., 1992). The laminar timing of border ownership may thus give clues regarding the roles of these pathways in this computation. Here, we used linear multielectrode probes to compare onset times of border ownership selectivity between laminar compartments.

It is also unknown how the preferences of border ownership-selective neurons in different layers relate to each other. This is in contrast to orientation preference, which is well known to be spatially organized in the primate brain. Primate areas V1 and V2 contain orientation columns (Hubel and Livingstone, 1987; Hubel et al., 1978; Vanduffel et al., 2002). In V4, imaging studies indicate that there is at least clustering of orientation preference in superficial layers of V4 (Ghose and Ts’o, 1997; Roe et al., 2012; Tanigawa et al., 2010), although it is unclear whether these clusters are columnar. Nor do we know if border ownership preference is organized in columns. The border ownership of a given border is defined entirely by asymmetries outside the cRF (as opposed to the border’s orientation). A systematic organization of border ownership preference would therefore imply a clustered arrangement of the neural pathways that underlie these asymmetries.

Finally, the relation between border ownership selectivity and orientation tuning is unclear. Prior studies have focused on orientation-selective units and tested border ownership at the neuron’s preferred orientation, without examining the relationship between orientation preference and border ownership preference (Hesse and Tsao, 2016; Zhang and von der Heydt, 2010; Zhou et al., 2000). One possibility is that border ownership is fundamentally a border property, such that border ownership selectivity is maximal for the preferred border orientation. Another possibility is that border ownership selectivity rather represents a surface signal (Grossberg, 2016), and may thus be less strictly tied to border orientation and orientation tuning.

Here we addressed these questions using laminar multielectrode probes to record from border ownership selective units in macaque area V4 across layers. We replaced the native dura with a transparent artificial dura to enable us to reliably position the probe normal to the cortical surface, on the relatively narrow exposed surface of area V4. We compared the timing and magnitude of border ownership selectivity across laminar compartments. If border ownership selectivity in V4 is inherited from V2, its dominant source of cortical input (Markov et al., 2011), we expect to see it early and prominently in neurons in the granular layer, which is the main target of this projection (Rockland, 1992; Gattass et al., 1997). Next, we compared the functional organization of border ownership preference across layers, to test whether the preferred side of ownership is shared. Finally, we examined the relationship between border ownership preference and border orientation preference.

## Results

To elucidate the functional organization and timing of border ownership we used laminar recording electrodes oriented normal to the cortical surface to record well-isolated units and multi-unit activity in macaque area V4, during fixation. We present data from two rhesus macaques recorded over 88 penetrations (animal Z: 34; animal D: 54). For each penetration, we first obtained data to map the receptive fields by presenting fast sequences of Gaussian windowed luminance contrast edges and rings at random positions in the appropriate visual quadrant. The orthogonal position of the probe relative to the cortical surface resulted in a vertical stacking of the receptive fields from different electrode contacts (Figure 1H,I; Figure 1-figure supplement 1C). At this retinotopic location we then obtained data to determine orientation tuning (luminance contrast edges of different orientations) and to compute current source density (see below). We then presented the border ownership stimuli (Figures 1A,C). These stimuli were similar to those that have previously been used to measure border ownership tuning (Zhou et al., 2000). We carefully positioned square stimuli such that one edge (termed the ‘central edge’) fell within the cRF, while ensuring that the stimulus features that defined border ownership (the other three edges and the four corners of the square) all fell outside of the cRF. Units for which the stimulus was not properly placed relative to the cRF were not included in the analysis (see inclusion criteria in Methods). This resulted in 685 well-isolated units (animal Z: 227; animal D: 458) and 765 multiunit clusters (animal Z: 329; animal D: 436).

Figure 1B shows the responses from a well-isolated unit to the stimuli in Figure 1A as raster plots. The unit fires more to stimulus 1 than to stimulus 2, even though the stimulus information in the cRF is identical in both cases. In other words, this unit fires more to an identical contrast edge in the cRF if that edge is owned by an object on the lower left compared to when it is owned by an object on the upper right. To test whether the difference in spike rates could be explained by the difference in luminance of the square object between stimulus 1 and stimulus 2, we also reversed all luminances (resulting in stimuli 3 and 4 in Figure 1A). Again we observe that this unit prefers the stimulus where the central edge is owned by an object on the lower left (prefers stimulus 3 over stimulus 4, Figure 1B). Together, we conclude that this unit’s preferred side of border ownership is the lower left side of the border, indicated by the direction of the green arrow in Figure 1B. We assessed statistical significance of border ownership selectivity with a permutation test on the absolute value of the border ownership index (*BOI*, see Methods, |*BOI|* = 0.38 for this unit, permutation test p<0.0001).

To estimate the laminar position of units, we performed current sourced density (CSD) analysis of the local field potential evoked by small rings in the cRF (Methods; Figure 1-figure supplement 1; Nandy et al., 2017; Mitzdorf, 1985). This analysis results in a pattern of current sinks and current sources (Figure 1F shows an example CSD map for the penetration during which the unit in Figure 1B was recorded). Such maps show a prominent leading current sink in the central portion of the penetration (white star in Figure 1F), with a current source followed by a current sink on the electrode contacts positioned deep from it (white disc in Figure 1F). Below this current source, we typically observe a current sink with longer latency (white triangle in Figure 1F). This sink-source pattern occurred consistently in different penetrations (Figure 1G; Figure 1-figure supplement 1; Figure 1-figure supplement 2). Studies from other laboratories in behaving macaques have described this sink-source pattern as well in extrastriate areas, not only in area V4 (Pettine et al., 2019, their Figure 1C; Lu et al., 2018, their Figure S4B) but also downstream in the medial temporal cortex (Takeuchi et al., 2011, area 36, their Figure S1, note that the ordinate is reversed in this figure relative to ours). Takeuchi et al. paired this analysis with histological verification and found that the prominent early current sink indicates the position of the granular layer. We therefore draw boundaries above and below this current sink and identify this compartment as the granular layer (white star, between white dashed and dotted lines in Figure 1F). From the position of the electrode contact on which each unit was recorded relative to the CSD map, we could then locate units in superficial, granular or deep layer compartments (see Methods for classification criteria). For example, the unit shown in Figure 1B was located in the deep compartment. Figures 1C,D,G,I show another example (multiunit cluster), recorded in deep cortical layers from a different penetration (|*BOI|* = 0.51, p<0.0001).

Across the population, we find border ownership selectivity in 51.1% of well-isolated units (350 out of 685 units; pooling across well-isolated and multiunit clusters, we find border ownership selectivity in 44.6% of units (647 out of 1450 units)). This proportion is high in all laminar compartments (superficial: 43.3% (58 out of 134); granular: 57.3% (102 out of 178); deep: 56.0% (121 out of 216); pooled well-isolated and multiunit clusters: superficial: 42.8% (140 out of 327); granular: 51.5% (167 out of 324); deep: 49.1% (210 out of 428)).

### Border ownership selectivity occurs first in deep cortical layers

Figure 2A shows the time course of the responses evoked by the preferred (solid red line) and non-preferred (dashed blue line) side of border ownership, averaged over all well-isolated units that are selective for border ownership. Consistent with prior studies (Zhou et al., 2000), we observe that border ownership selectivity (difference in response to the preferred and the non-preferred side) occurs early after onset of the stimulus-evoked response. The asterisk indicates when the difference between these functions first becomes statistically significant (56.5 ms after stimulus onset; sign rank test p<0.05 for 20 adjacent ms). When evaluating these functions in each laminar compartment we observe that these functions diverge substantially early after onset in deep cortical layers (Figure 2D; significant at 58.8 ms), but later in superficial (Figure 2B; significant at 85.8 ms) and granular layers (Figure 2C; significant at 67.3 ms). This definition of latency depends on sample size, but a subsampling analysis shows that differences in sample size between layers do not explain the shorter latency for deep layers (Figure 2-figure supplement 1). To compare the time course of border ownership modulation between layers, we defined the border ownership index function *B* for each laminar compartment as

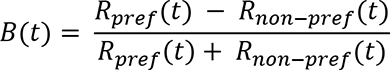

where *R*_*pref*_(*t*) and *R*_*non*−*pref*_(*t*) are respectively the evoked response functions to the preferred and the non-preferred sides of border ownership (red and blue dashed functions plotted in Figure 2A-D). *B*(*t*) is plotted for each laminar compartment in Figure 2E, confirming that border ownership modulation rises earlier in the deep layers than in the other laminar compartments. We quantified the difference by defining latencies on these functions as the crossing with a threshold defined from the null distribution obtained by shuffling the stimulus labels (see Methods). Latency was significantly shorter for deep layer units (75.8 ms, 95%CI [68.4 85.2]) than for granular layer units (94.7 ms, 95%CI [82.2 105.7], bootstrap procedure (see Methods) p=0.006) and for superficial layer units (97.7 ms, 95%CI [78.0 103.7], bootstrap procedure p=0.018). The same was true when well-isolated units and multi-units were pooled (deep: 78.2 ms, 95%CI [73.3 88.2], n=210; granular: 94.0 ms, 95%CI [87.9 106.5], n=167; superficial: 100.7 ms, 95%CI [85.1 104.6], n=140; deep vs. granular: p=0.007; deep vs. superficial: p=0.009). To verify the robustness of the temporal differences between layers, we also evaluated the time course of border ownership selectivity using another method, by evaluating border ownership reliability (Figure 2F; introduced by Zhou et al., 2000; detailed in Methods). Briefly, this metric quantifies the reliability of border ownership tuning when comparing spike counts between single trials to stimuli with opposite border ownership. Reliability values correspond to the proportion of such single trial comparisons for which the spike count is highest for the border ownership condition that is preferred across trials. We computed border ownership reliability in 100 ms-sliding windows (Figure 2F; latency defined similarly as in Figure 2E, using right edge of the analysis window). Again we find that border ownership reliability rises earlier in the deep layers (89.9 ms, 95%CI [81.5 99.9]) than in the granular (105.4 ms, 95%CI [94.9 114.6]; bootstrap procedure p=0.015) and superficial layers (109.4, 95%CI [98.9 124.9]; p=0.006). After border ownership selectivity has been established, the average border ownership index tends to be higher in the deep compartment (Figure 2E; border ownership index between 200 and 500 ms after stimulus onset: mean ± s.e.m. 0.50 ± 0.02) than in the granular (0.43 ± 0.02) and superficial compartment (0.45 ± 0.03), but the differences do not reach statistical significance (deep vs. granular: Wilcoxon rank sum test p=0.051; deep vs. superficial: p=0.22). Border ownership reliability saturates around 0.85 in all three compartments (Figure 2F). Together, these data indicate that border ownership selectivity does not occur first in granular layer units but instead in deep layer units.

**Figure 2.**
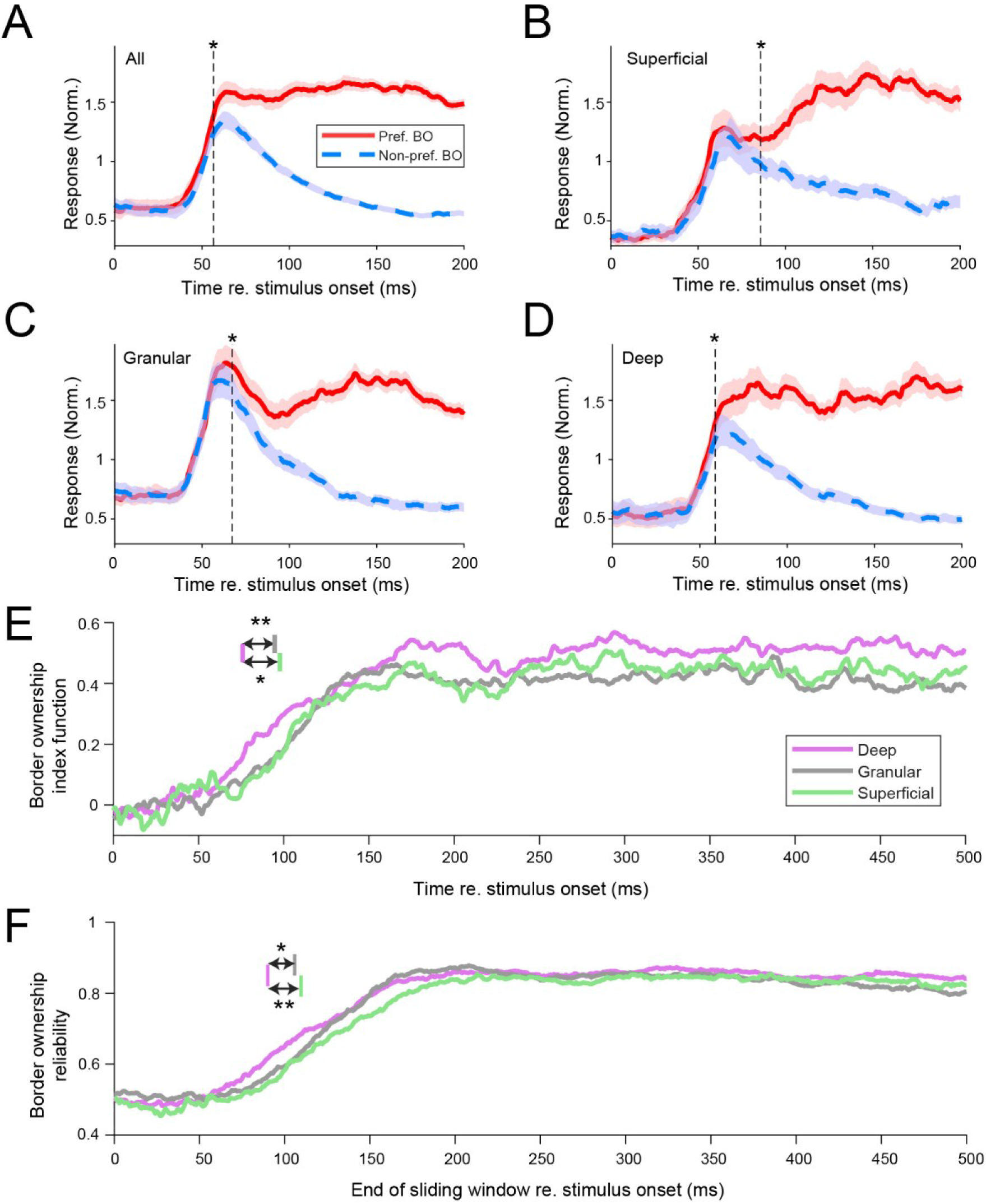
Border ownership selectivity occurs first in deep cortical layers. (**A**) Response time courses for the population of border ownership selective well-isolated units (n = 350). Functions are plotted separately for the responses to the preferred side of border ownership (solid red line; mean +/- s.e.m.) and the non-preferred side of border ownership (dashed blue line). The vertical dashed line indicates when the difference between the functions first becomes significant (Wilcoxon signed rank test, p<0.05 for ≥20 ms). (**B-D**) Similar to A, for the subpopulations of well-isolated units selective for border ownership that could be located respectively to superficial (B), granular (C) and deep (D) layers (see Methods for criteria of layer assignment). Superficial: n = 58 units; granular: n = 102 units; deep: n = 121 units. Analysis as in panel A. See also Figure 2-figure supplement 1. (**E**) Border ownership index functions of the different laminar compartments (see Methods). Colored vertical lines indicate latency for the different layers, defined as the earliest crossing of the border ownership response function for ≥20 ms with the threshold. The threshold was set at the level for which <1% of functions obtained after shuffling the stimulus labels had a defined latency (0.156). Latency, deep: 75.8 ms, 95%CI [68.4 85.2]; granular: 94.7 ms, 95%CI [82.2 105.7]; superficial: 97.7 ms, 95%CI [78.0 103.7]. **: bootstrap procedure (see Methods) p=0.006; *: p=0.018. See also Figure 2-figure supplement 2. (**F**) Border ownership reliability calculated in a 100-ms sliding window for the three laminar compartments. Colors as in E. Colored lines in top of panel indicate the earliest crossing for ≥20 adjacent ms with the threshold, defined similarly as for panel E. Threshold crossings: deep 89.9 ms, 95%CI [81.5 99.9]; granular 105.4 ms, 95%CI [94.9 114.6]; superficial 109.4 ms, 95%CI [98.9 124.9]. **: bootstrap procedure p=0.006; *: p=0.015.

To test whether the short latency in deep layers is specific for border ownership or a general feature of this laminar circuit, we performed two additional analyses. First, we evaluated the latency of spikes evoked by small ring stimuli in the cRF. For these responses we find, in contrast to border ownership selectivity, that the latency is shorter in the granular layer compared to the deep layer and superficial layers (Figure 3; latency defined as crossing of the functions with a threshold value halfway between baseline and peak). Note that these responses are derived from the same stimuli used to compute the CSD maps, but represent a different signal (spiking responses as opposed to the current sink-source patterns from local field potentials used to define the laminar compartments). Second, we evaluated the responses to the border ownership stimuli for another type of selectivity, contrast polarity. This refers to the relative luminance contrast across the edge, i.e. the difference between panel 1 and panel 3 (or between panel 2 and panel 4) in Figure 1A. For the contrast polarity index functions, we do not find an earlier rise in selectivity in the deep layers, even though they are derived from the same units as in Figure 2E (Figure 2 – figure supplement 2). Together, these data indicate that the earlier selectivity in deep layers compared to granular and superficial layers is specific for border ownership.

**Figure 3.**
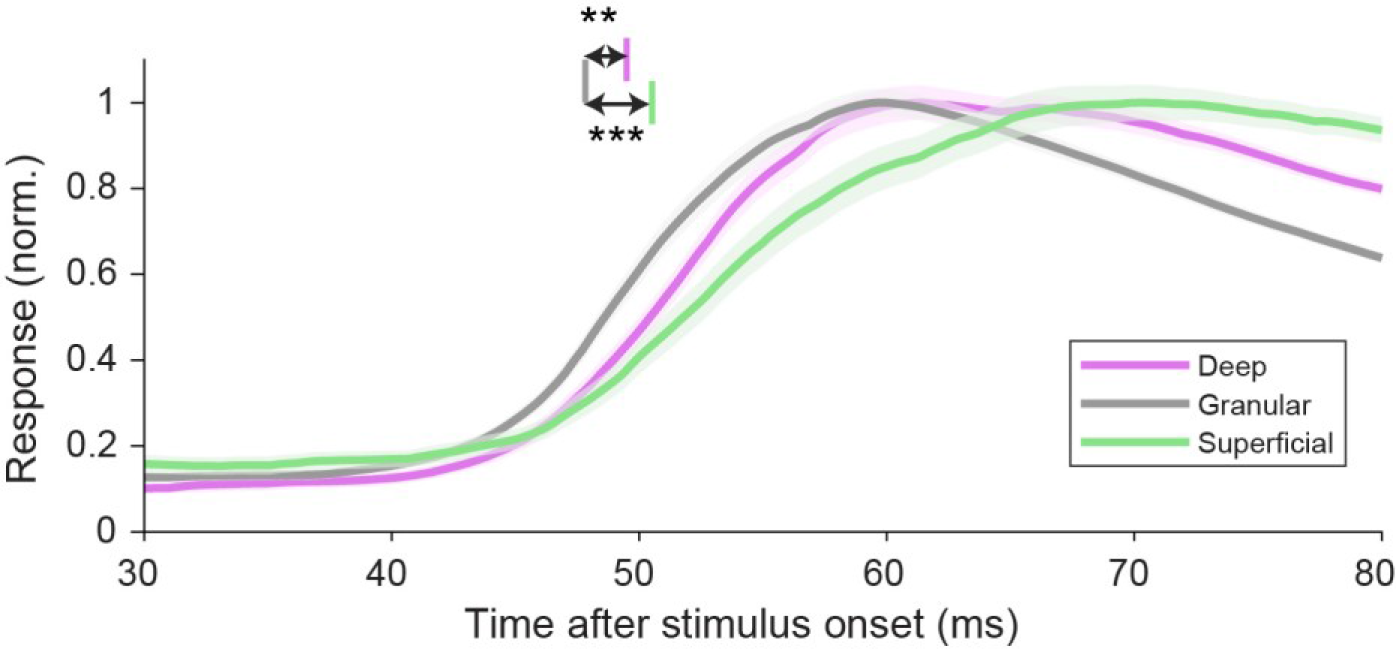
Spike latencies evoked by flashed stimuli appearing in the classical receptive field are shortest in the granular layer. Response time courses of spikes evoked by small rings centered on the classical receptive field, for well-isolated units recorded in each laminar compartment (mean ± s.e.m.). Vertical lines indicate mean latencies for each laminar compartment (defined as crossing of the functions with a threshold value halfway between baseline and peak, 0.435). Deep: 49.4 ms, 95%CI [48.3 50.3], 173 units; granular: 47.8 ms, 95%CI [47.2 48.5], 152 units; superficial: 50.5 ms, 95%CI [49.1 52.2], 101 units. ***: bootstrap procedure p=0.0005; **: p=0.002.

### The preferred side of border ownership is organized in columnar clusters

How does the preferred side of border ownership compare between units recorded in a column of cortex? For a given edge, for example vertical, there are two possibilities for preferred side of border ownership: left or right from the edge. Border ownership for such an edge has been assumed to be represented by the activity of two oppositely tuned subpopulations (green and purple in Figure 4A, top; Craft et al., 2007). Our data show indeed that these two subpopulations exist in similar proportions (Figure 4A, top). The same is true for units that encode border ownership for horizontal edges (Figure 4A, bottom), and when we express the preferred side of border ownership not relative to the screen, but relative to the fixation point (Figure 4–figure supplement 1). This indicates that for a given edge, the two possible sides of border ownership are encoded by distinct subpopulations of neurons that are similar in size.

**Figure 4.**
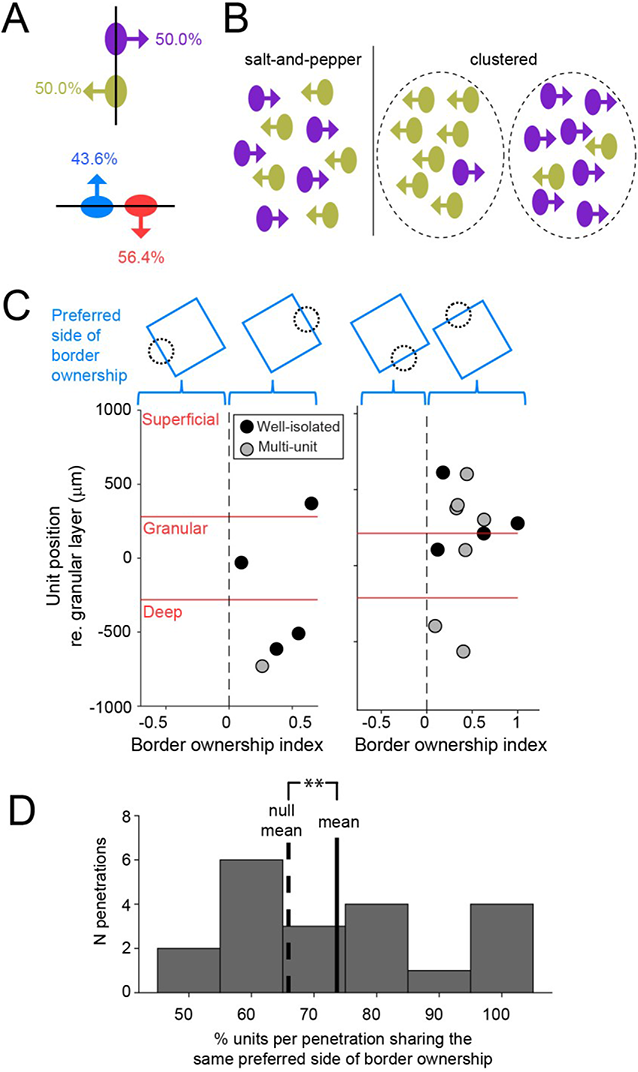
Preferred side of border ownership is clustered. (**A**) For a given edge, there are two possibilities for preferred side of border ownership. Percentages indicate the fraction of well-isolated units selective for border ownership that have the preferred side of border ownership indicated by the arrow (vertical edge: n = 64; horizontal edge: n = 78). See also Figure 4-figure supplement 1. (**B**) Cartoon showing possibilities for spatial organization of neurons with opposite preferred sides of border ownership, for a vertical edge. (**C**) Each panel shows border ownership selective units recorded during one penetration. The abscissa shows the signed *BOI* of each unit (sign indicates preferred side of border ownership, shown above the panels). Ordinate shows the position of the units relative to the center of the granular layer. (**D**) Histogram of the proportion of units per penetration that shares the same preferred side of border ownership, for all penetrations with at least four border ownership-selective well-isolated and multiunits spanning from superficial to deep cortical layers. Solid line is distribution mean, dashed line shows null distribution mean. **: p=0.003 (20 penetrations with 136 units).

These two subpopulations could be mixed in a salt-and-pepper pattern, or they could be clustered according to their preferred side of border ownership (Figure 4B). Figure 4C shows the signed *BOI* for all well-isolated units and multiunit clusters selective for border ownership for two example penetrations. The sign of *BOI* indicates which side of the border is preferred by each unit, as indicated by the cartoons above the panels. For both of these penetrations, all border ownership units recorded on the same probe prefer the same side of border ownership. In the population of all penetrations with at least four border ownership units, the proportion of units that share the same preferred side of border ownership is significantly higher than chance (well-isolated units: 74.4%, randomization test p=0.017, 23 penetrations with 110 units; including multiunit clusters: 71.7%, p=0.005, 47 penetrations with 288 units). Also in the subgroup of penetrations with border ownership units spanning from superficial to deep layers, we find significant clustering (Figure 4D; well-isolated units: 81.1%, randomization test p=0.010, 6 penetrations with 30 units; including multiunit clusters: 73.7%, p=0.003, 20 penetrations with 136 units). Together, these data indicate that border ownership preference is organized in spatial clusters that span cortical layers in a columnar fashion.

### Columnar organization of orientation selectivity

Border ownership acts on edges that are oriented. Similar to our finding that preferred border ownership is clustered, several imaging studies have shown that preferred orientation is spatially clustered in V4 in domains (e.g. Li et al., 2013; Tanigawa et al., 2010), but it is not known whether these form columns. To test whether orientation preference in V4 is shared in clusters that extend vertically across layers, we determined orientation tuning for the units in our sample from responses to luminance contrast edges centered on the cRF (independent data set from the border ownership stimuli, see Methods). Our data confirm clustering of preferred orientation and reveal that these clusters span across laminar compartments in V4 (Figure 5). Each polar plot in Figure 5 shows a penetration with at least four orientation-selective well-isolated units or multiunit clusters. Solid vectors indicate the preferred orientation of each orientation-selective unit as the resultant vector of the responses to edges of different orientations, and color indicates laminar compartment (open symbols indicate multiunit clusters). The preferred orientation for a penetration was then calculated as the resultant vector across the vectors of all orientation-selective units in that penetration, and its angle is indicated by the blue dashed line. The significance of clustering of orientation preference was assessed by comparing the magnitude of this resultant vector against a null distribution generated by randomizing the preferred orientation for each orientation-selective unit in the penetration and performing the same calculation. For each of these six penetrations, this resultant was significantly larger than expected from the null distribution (p < 0.05). This was true for 9 out of 18 penetrations with at least four orientation-selective well-isolated units distributed over all three laminar compartments (for 17 out of 34 penetrations when well-isolated units and multiunit clusters were pooled). These data suggest that orientation domains in V4 are columnar.

**Figure 5.**
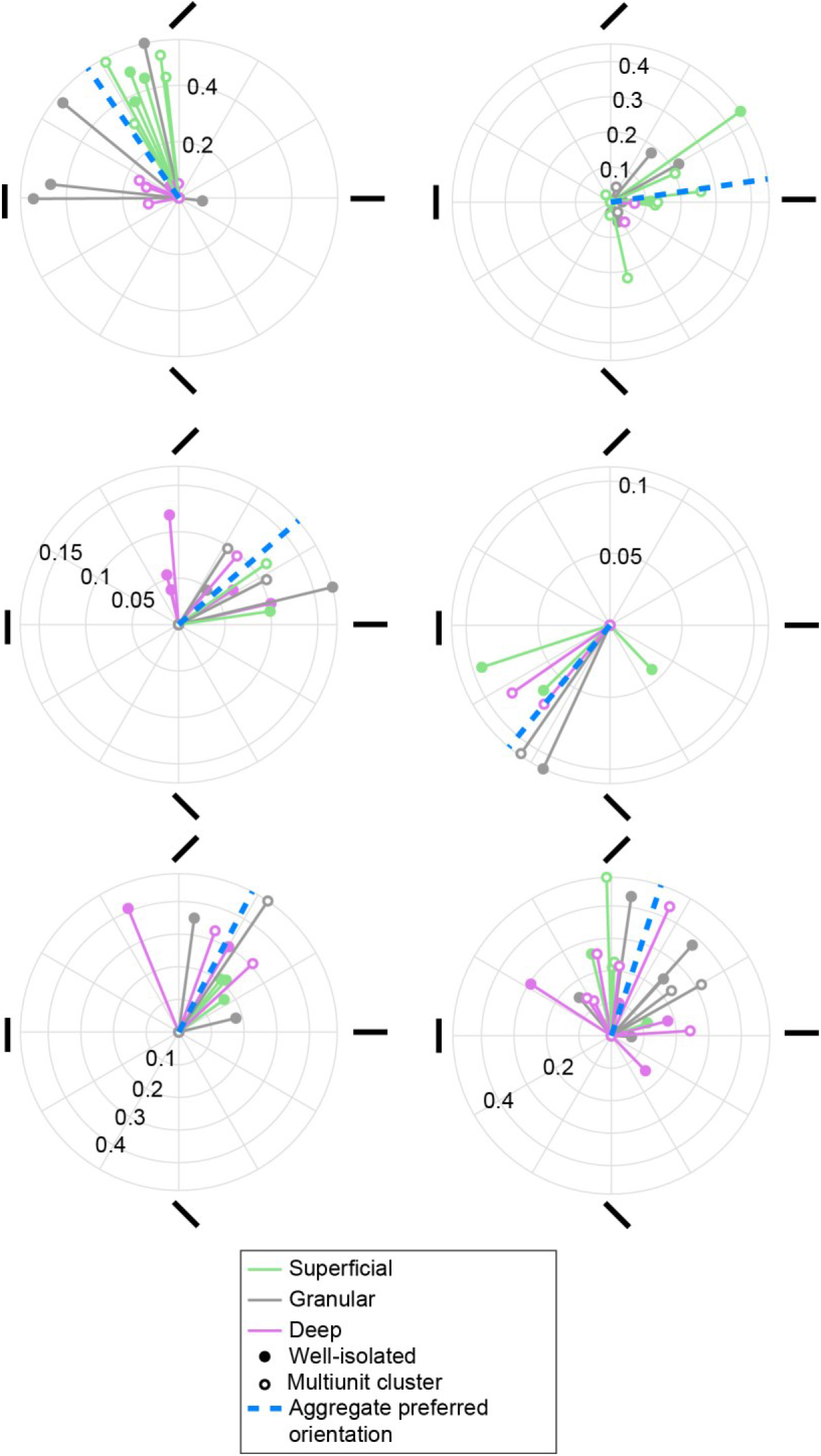
Columnar clusters of preferred orientation in V4. Polar plots are shown for six penetrations with at least four orientation-selective units distributed across all three laminar compartments. Each solid vector corresponds to one unit and indicates the preferred orientation (direction) and the degree of orientation tuning (magnitude), quantified respectively as the direction and magnitude of the resultant vector of the responses to each orientation. Color indicates the laminar compartment. Open symbols indicate multiunits, closed symbols indicate well-isolated units. Blue dashed line indicates the preferred orientation across units in a penetration (aggregate preferred orientation). All six penetrations shown have a significant aggregate preferred orientation.

### Border ownership selectivity often occurs far away from the preferred orientation

Having found that both border ownership preference and orientation preference are organized as columnar clusters in V4, we next asked what the relation was between orientation selectivity and border ownership selectivity. We find that border ownership selectivity is significantly more common in units that are selective for orientation than in those that are not, but it is also surprisingly common in units that are not selective for orientation (Table, respectively 53.3% and 36.8%, Chi square = 9.51, p=0.002; including multiunit clusters: respectively 45.9% and 31.3%, Chi square = 19.5, p=0.00001).

**Table.**
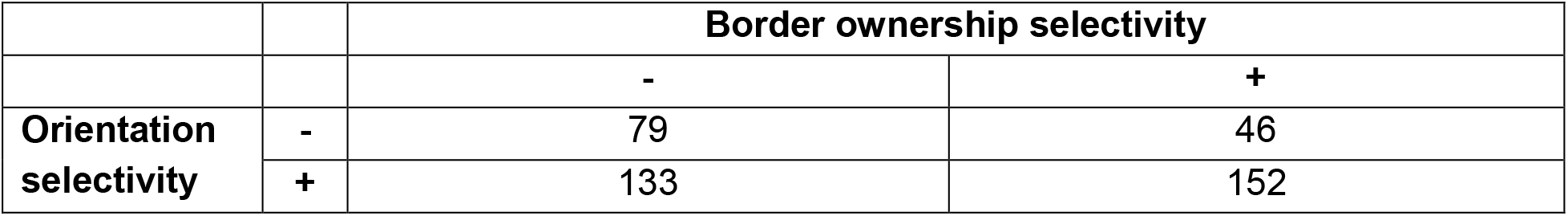
Border ownership selectivity is more common in orientation-selective well-isolated units but often occurs in units that are not selective for orientation.

Three example orientation-selective well-isolated units are shown in Figure 6A. These polar plots are organized according to the location of the square object relative to the cRF, for different orientations of the central edge (the edge that runs through the cRF). This is indicated by the stimulus cartoons around the plot. For opposite locations on the polar plot, the central edge has thus the same orientation, but is owned by a square positioned on opposite sides of the cRF (opposite border ownership). The *BOI* for each tested orientation of the central edge is shown by a red vector (filled red circles indicate statistically significant border ownership selectivity). The amplitude of this vector corresponds to |*BOI*|, and the polarity of the vector points towards the square location that corresponds to the preferred side of border ownership for that orientation of the central edge. For example, in case of a vertical central edge, the third unit (cyan triangle) prefers that that edge is owned by a square on the right (if it would have preferred the vertical central edge to be owned by a square on the left, the red vector that points down would have pointed up). The black line indicates the preferred orientation of a luminance contrast edge in the cRF for each unit. For the left unit in Figure 6A (red star), this preferred edge orientation matches with the orientation of the central edge for which border ownership selectivity is significant (filled red circle). This is the expected scenario: border ownership has been assumed to act on edges at the preferred orientation, and prior studies have used edges at the preferred orientation to test border ownership selectivity (Zhou et al., 2000). At the population level, the average orientation of edges with border ownership selectivity is biased towards the preferred orientation for edges (visible as the clustering of the data points around the identity line in Figure 6C; average distance 27.6°, randomization test (see Materials and methods) p=0.018; n= 68 well-isolated units tested with at least four orientations); distance distribution shown in Figure 6D; including multiunit clusters: p=0.017; n=159). Note that both plotted variables in Figure 6C are periodic and two periods are shown for each (thus each data point is plotted four times; the grey square outlines an area corresponding to one period for both variables). Orientation-tuned units thus tend to be border-ownership selective for edges with an orientation near their preferred orientation.

**Figure 6.**
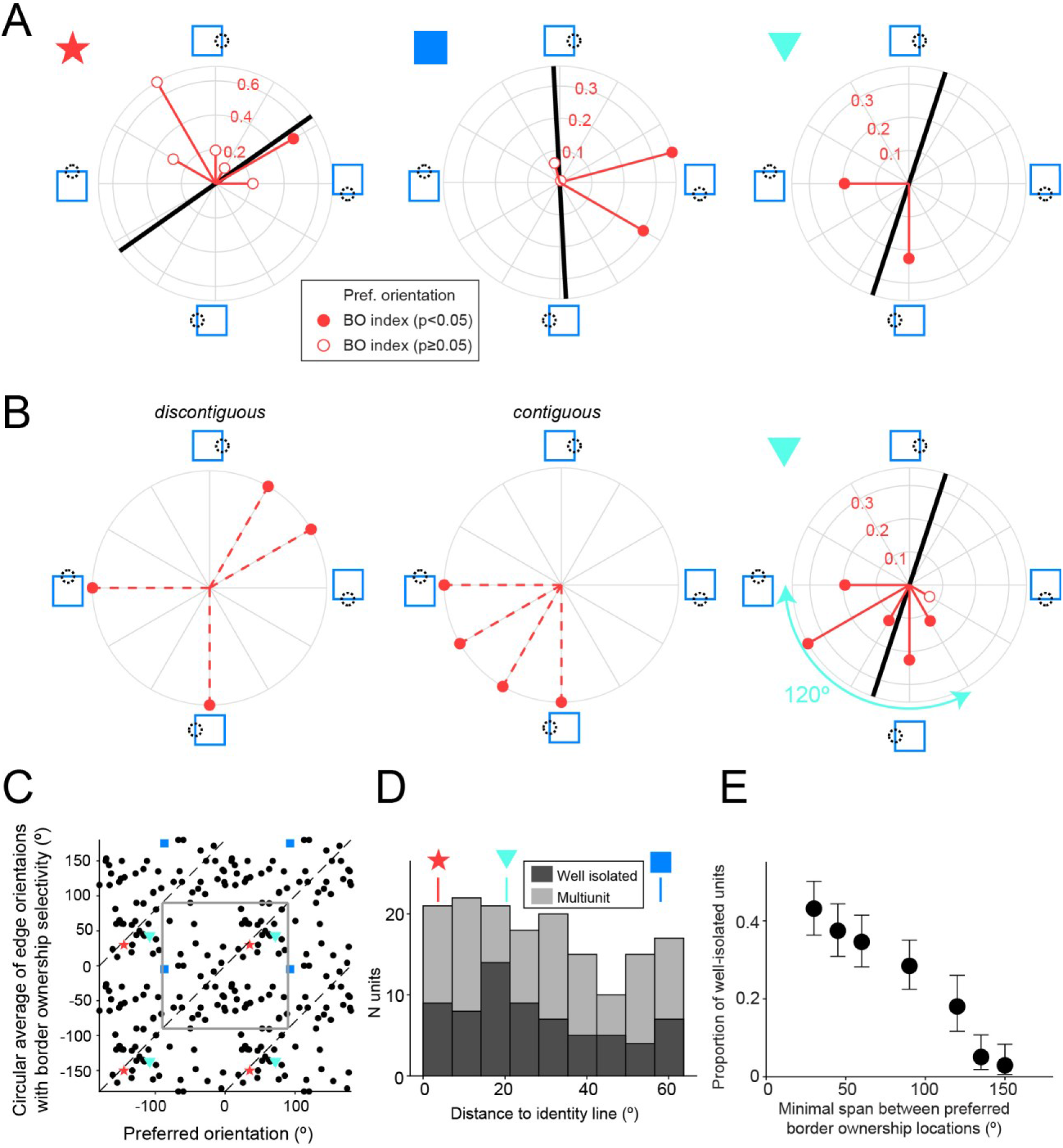
Border ownership selectivity often occurs far away from the preferred orientation. (**A**) Polar plots for three example units that were tested for border ownership selectivity with edges of multiple orientations (orientations of red vectors indicate tested orientations). Polarity of red vector relative to origin indicates the preferred side of border ownership for each edge orientation, according to the stimulus cartoons surrounding the plot, and filled circles indicate cases with statistically significant border ownership selectivity (adjusted for multiple comparisons using Bonferroni correction). Magnitude of red vectors corresponds to |*BOI*|. Black line indicates preferred edge orientation. Colored symbols on the upper left side of each polar plot correspond to the symbols in C-D. See also Figure 6-figure supplement 1. (**B**) Left and middle: for units showing border ownership selectivity to orthogonal edges there are two possibilities for the intermediate edge orientations. Either the preferred sides of border ownership are spatially discontiguous with those for the orthogonal edges (left), or they are contiguous (middle). Right: polar plot for the same unit as in A (right polar plot), now including data for the orientations that were not shown in A. (**C**) Scatter plot comparing the preferred orientation for edges (abscissa) with the circular mean of edge orientations for which border ownership selectivity is significant (ordinate), for all units that were tested for border ownership for at least four orientations (n = 68 well-isolated units). Note that each data point is plotted four times, because both variables are periodic and two periods are shown for each. The grey square outlines an area corresponding to one period for both variables. Dashed lines indicate identical values. Colored symbols correspond to the units in A,B,D. (**D**) Smallest orthogonal distance to the identity line for the units shown in C. The data is significantly biased towards zero (see Results). Distance values for the example units shown by colored symbols in A-C are indicated above the histogram. (**E**) Proportion of border ownership well-isolated units where the preferred sides of border ownership cover an angular span at least as wide as the value indicated along the abscissa (for all units tested with such spans). For example, the unit in B (right) has a span of preferred sides of border ownership of 120°. Note that the span between spatially contiguous preferred sides of border ownership is necessarily ≤180°. N units for each value of on the abscissa: ≥30°: 211; ≥45°: 211; ≥60°: 211; ≥90°: 211; ≥120°: 122; ≥135°: 118; ≥150°: 102.

That being said, the scatter in Figure 6C is substantial. Some units only have border ownership selectivity for orientations that are far from the identity line, such as the unit indicated with the blue square in Figure 6A (middle panel, corresponding to the same symbol in Figure 6C). This unit shows, paradoxically, significant border ownership selectivity for edge orientations that are nearly orthogonal to the preferred edge orientation. This is true for two edge orientations (filled red circles), suggesting that this misalignment is systematic. We ruled out that this is related to a difference in orientation preference for isolated edges versus edges that are part of squares by comparing orientation tuning for both data sets. Figure 6–figure supplement 1A shows that the preferred orientations for both stimuli match very well for this unit (black line vs cyan line), and this is true for the population as well (Figure 6–figure supplement 1B).

Furthermore, we find that border ownership selective units often show border ownership selectivity to edges that maximally differ in orientation, i.e. orthogonal edges. An example unit is shown in the right panel in Figure 6A (cyan triangle). This is true for 29.7% of border ownership selective well-isolated units (n=182; 24.4% including multiunits, n=353) that were tested with sets of squares at orthogonal angles. In such cases, the preferred side of border ownership for a central edge with an orientation in between those orthogonal orientations could in theory be spatially discontiguous (Figure 6B, left panel) or contiguous (Figure 6B, middle panel) with the preferred sides of border ownership for the pair of orthogonal orientations. We find that for all 24 out of 24 well-isolated units that were selective for the border ownership of squares at orthogonal and intermediate orientations, the preferred side of border ownership for these intermediate orientations was spatially contiguous (such as the example in Figure 6B, right panel; including multiunits, this is true for 35 out of 35 units). Those units thus have a wide but spatially contiguous area of preferred sides of border ownership. For example, the unit in Figure 6B (right panel) has an area of preferred sides of border ownership that spans 120° (cyan double arrow). A span of at least this width occurs for 18.0% of border ownership units (Figure 6E; 15.4% including multiunit clusters).

## Discussion

The correct assignment of borders is a key step in the process of constructive perception that enables us to identify and localize objects in a visual scene. Prior studies have found neurons that signal border ownership, but the underlying circuitry is not understood. Various theoretical accounts have been offered. One group of models hypothesizes that border ownership selectivity arises in a feedforward manner, through the successive elaboration of progressively more complex receptive field properties (Sakai et al., 2012; Sakai and Nishimura, 2006; Supèr et al., 2010). Other accounts posit that border ownership assignment crucially depends on horizontal- and cortico-cortical feedback signaling (Craft et al., 2007; Grossberg, 2016; von der Heydt, 2015; Zhang and von der Heydt, 2010; Zhaoping, 2005). These different pathways differ in their laminar organization. Feedforward inputs typically terminate dominantly in the granular layer, whereas long-range horizontal fibers and cortical feedback projections avoid the granular layer (Douglas and Martin, 2004; Rockland and Pandya, 1979; Rockland et al., 1994; Rockland and Lund, 1983; Ungerleider and Desimone, 1986; Yoshioka et al., 1992).

### Border ownership selectivity and the laminar circuit

Given that about half of the neurons responsive to edges in V2 are selective for border ownership (Zhou et al., 2000), neurons in V4 could simply inherit this property from V2 through the feedforward projection from V2 to V4. This projection is the most important source of feedforward input to V4 (Markov et al., 2011), and targets mostly layer 4 (granular layer) in V4 (Gattass et al., 1997; Rockland, 1992). Our data shows that spike latency for stimuli in the classical receptive field is shortest in the granular layer, which is similar to what has been described in other areas in macaque visual cortex (V1: e.g. Nowak et al., 1995; MT: Raiguel et al., 1999), and consistent with work by others in awake V4 (Lu et al., 2018). Note that this may not be universal across areas: in anesthetized V2 spike latency has been reported to be longer in layer 4 than in infragranular layers (Nowak et al., 1995).

If this feedforward projection would thus simply provide the earliest border ownership signals in V4, we would thus detect border ownership selectivity in the granular layer at least as early as in the other compartments, or earlier. While we do observe such a pattern for responses evoked by flashed stimuli appearing within the classical receptive field, and also for the emergence of selectivity for contrast polarity (Figure 2-figure supplement 2), we do not observe this pattern for selectivity to border ownership. Instead, we find that deep layer neurons compute border ownership selectivity significantly earlier than neurons in the granular layer and in the superficial layers. One might argue that these data are consistent with V2 afferents preferentially targeting apical dendrites of deep layer neurons that extend in the input layer. But in that case, we would expect shorter spike latencies for deep layer neurons than for granular layer neurons irrespective of the cue or stimulus. This is, again, not what we find: spike latencies for responses evoked by small stimuli in the cRF are shorter in the granular layer than in the deep layers, and also for contrast polarity we do not observe the earliest selectivity in deep layers in V4.

Another possibility is that deep layer neurons perform this computation earliest solely relying on an integration of feedforward signals from the V2-to-V4 projection relayed by different granular layer neurons. We think this is unlikely. First, this would mean that those relaying granular layer neurons do not receive sufficient contextual information to become selective for border ownership as fast as the deep layer neurons that integrate their output, despite the substantial input that granular layer neurons receive from other granular layer neurons (Xu et al., 2016). Second, while deep layer neurons certainly receive input from granular layer neurons, the densest projections from granular layer neurons typically target superficial layers (e.g. macaque V1: Lund and Boothe, 1975; Callaway, 1998; rat barrel cortex: Lübke et al., 2000). If that projection leads to the first computation of border ownership signals, one would thus expect to observe these signals in superficial layer neurons as fast as in deep layer neurons, in contrast with our data.

We think that the earlier computation of border ownership selectivity by neurons in deep layers is more likely a consequence of their unique properties. Scanning laser photo-stimulation studies in primary visual cortex showed that layer 5 neurons receive significant input from nearly all cortical layers, regardless of cell type, as opposed to neurons in other layers (Briggs and Callaway, 2005; Xu et al., 2016). Deep layers of the cortex, including in primates, include tall pyramidal cells whose apical dendrites reach up to layer 1, which could thus directly sample and integrate afferent information that arrives in a wide range of layers (Binzegger et al., 2004; Callaway, 1998; Lund and Boothe, 1975; Markov et al., 2014; Zarrinpar and Callaway, 2016). Such neurons thus seem well suited to integrate information provided by feedforward input with contextual information provided through corticocortical feedback and horizontal connections (Harris and Shepherd, 2015). Cortico-cortical feedback in V4 terminates densely in all layers except layer 4 (Markov et al., 2011; Maunsell and van Essen, 1983; Rockland et al., 1994). Intra-areal horizontal connections are prominent in layer 2/3 and in layer 5 (Lund et al., 1993; Yoshioka et al., 1992; Douglas and Martin, 2007). This may set these deep layer neurons up to be able to integrate the border in the classical receptive field (light grey arrow in Figure 7) with visual context from a wide region of space (dark grey arrows in Figure 7), which is required to compute border ownership selectivity. Indeed, studies in V2 showed that border ownership selectivity does not rely on a small number of localized object features but instead occurs through integration of extra-classical stimulus features over large areas of visual space (Zhang et al., 2010). Specialized intrinsic properties could further assist in this integration, such as the calcium spikes in apical dendrites of layer 5 neurons that can amplify the effects of feedback inputs (Takahashi et al., 2016).

**Figure 7.**
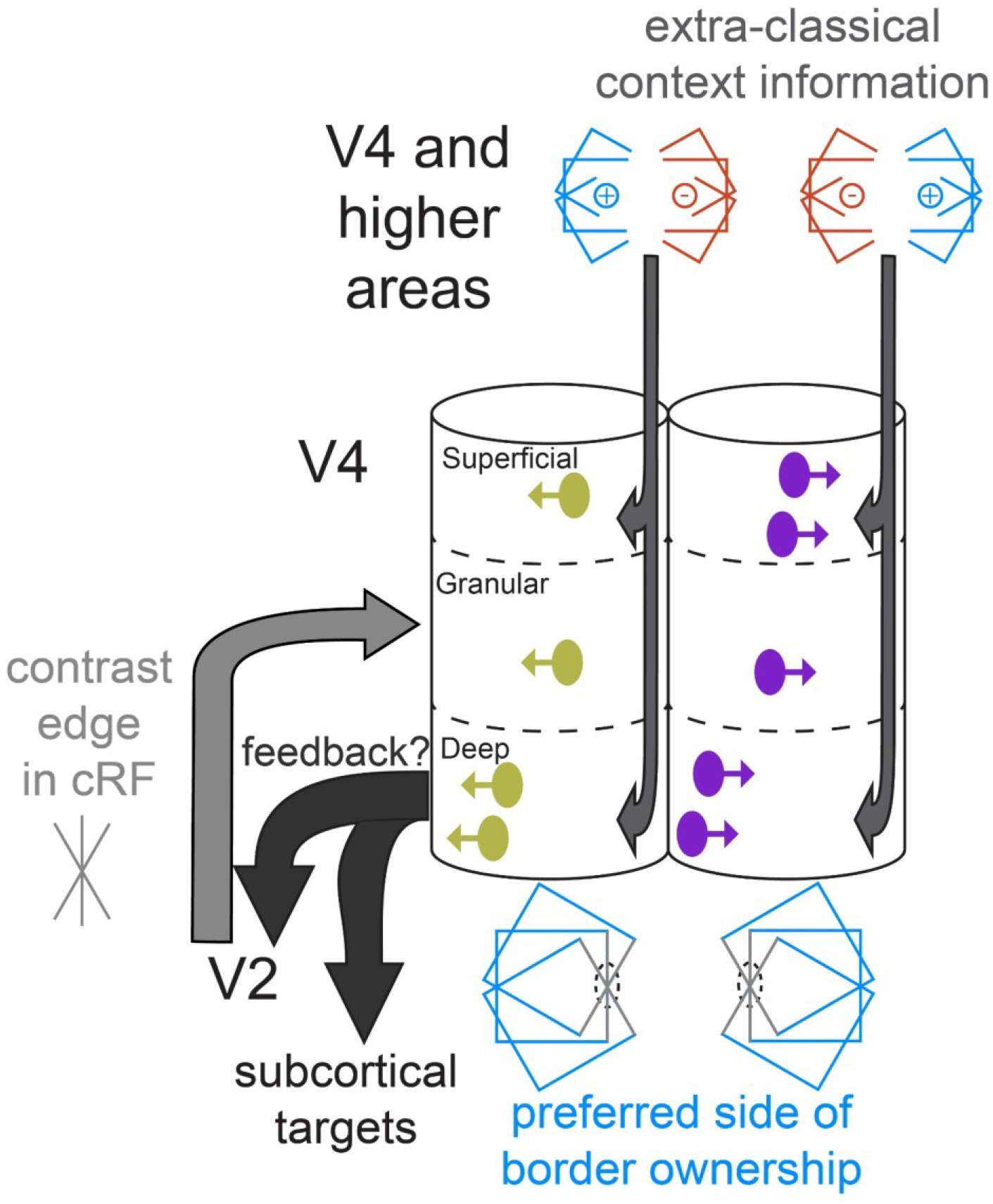
Cartoon of a conceptual model of columnar processing of border ownership, supported by our data. Two columnar structures are shown, each of which responds to vertical edges in the cRF, but they have opposite preferred sides of border ownership (indicated by the direction of the arrows on the units in the columns, and by the symbol below each column). Such columns may or may not be adjacent. The earliest occurrence of border ownership selectivity in the deep layers is symbolized by the positon of deep layer units near the left edge of the column. Light and dark grey arrows indicate where different types of stimulus information likely arrive first. Stimulus information in the cRF (grey edge) arrives first in the granular layer. Deep layer neurons may first compute border ownership selectivity by integrating this information with contextual information that arrives in superficial and deep layers, possibly provided by corticocortical feedback and horizontal connections. The content of these context signals is necessarily asymmetric to result in border ownership preference, and different for the two columns shown (indicated by the symbols above the columns). Because of their prominence in deep layers, early border ownership signals may be part of the feedback that projects from V4 to upstream areas (e.g. V2), and to subcortical targets, such as the superior colliculus (black arrows). Later on during the response, after border ownership signals have first been established in V4, the feedforward input from V2 to V4 likely also contains border ownership signals, since roughly half of V2 units are selective for border ownership (Zhou et al., 2000).

Our conclusion that the earliest border ownership signals in V4 are not inherited from V2 does not imply that the projection from V2 to V4 does not carry any border ownership signals. Indeed, since about half of V2 neurons are selective for border ownership (Zhou et al., 2000), this feedforward input most likely contributes to the border ownership signals in V4 later in the response. As argued in the next section, these border ownership signals in V2 may have been sculpted by border ownership-selective feedback from deep layers in V4.

Computational models have shown that myelinated feedback afferents carrying contextual information can indeed arrive fast enough to result in border ownership selectivity within ∼20 ms after response onset (Craft et al., 2007; Hu et al., 2019). An alternative mechanism based solely on horizontal fibers is unlikely, given the slow conduction velocity of these fibers (Girard et al., 2001; Zhang and von der Heydt, 2010), although, as discussed by Zhaoping (2005), it should be noted that data on horizontal fibers are scarce). That said, horizontal fibers could still provide part of the required contextual information: since these fibers are limited in length, their role could be to provide information from the portions of the stimulus closest to the cRF (reminiscent to their proposed role in surround suppression in V1 (Angelucci et al., 2017)). This would explain why the effect of near corners of squares tends to occur later than that of far corners of squares (Zhang and von der Heydt, 2010).

### Potential function of early border ownership signals in deep layers

Once border ownership selectivity has been established in V4 deep layers, these signals may contribute to visual processing along different pathways (black arrows in Figure 7). First, V4 is the dominant source of cortical feedback to V2, and 75% of these feedback neurons are located in deep layers in V4 (Markov et al., 2014). Early border ownership signals in deep layers of V4 may thus sculpt border ownership selectivity in V2. Second, V4 deep layers include neurons that project to the superior colliculus (Fries, 1984; Gattass et al., 2014). The superior colliculus may thus receive early border ownership signals from V4, which could contribute to planning upcoming saccades. Saccades target objects more frequently than background (Rothkopf et al., 2007). Early border ownership signals in deep layer neurons that project to the superior colliculus could serve to facilitate the rapid foveation of objects. Our data show that these signals are established in deep layers well before 100 ms after stimulus onset. They are thus computed quickly enough to allow time for saccade planning within the inter-saccadic interval, which is typically on the order of 200-300 ms (Otero-Millan et al., 2008). This proposed early role for deep layer neurons in supporting saccadic decisions is consistent with recent findings showing that deep layer V4 neurons encode more information about the direction of planned eye movements than do superficial neurons (Pettine et al., 2019). Border ownership signals may thus play a role beyond visual perception, perhaps including a role in guiding rapid oculomotor behavior.

### Border ownership and orientation tuning

Prior studies on border ownership only evaluated border ownership selectivity at the preferred orientation of each neuron (Hesse and Tsao, 2016; Zhang and von der Heydt, 2010; Zhou et al., 2000). In our data, there is indeed a bias towards border ownership selectivity for edges near the preferred orientation. However, we find that there is substantial scatter (Figure 6C,D). This breadth of tuning does not seem to stem from measurement noise. First, there is a good match between preferred orientation, as estimated from two separate data sets: one based on recordings made with isolated edges, the other using edges that are part of squares (Figure 6–figure supplement 1), suggesting that our estimates of orientation preferences are highly reliable, with relatively tight bounds. Second, for some units, border ownership selectivity, surprisingly, systematically only occurs for edges with orientations that are nearly orthogonal to the preferred orientation (Figure 6A, middle). Third, almost a third of units that are selective for border ownership to an edge with a given orientation are also border ownership selective to an edge that is orthogonal to that orientation. This reflects a systematic preference, because the preferred sides of border ownership in such cases form a single contiguous area in retinotopic space (Figure 6B). Finally, a substantial portion of units that are selective for border ownership are not selective for orientation (**Table**). Together, these data indicate that the relation between orientation and border ownership is much richer than has previously been appreciated. Encoding of orientation and of border ownership may represent separate axes of representation. This is not inconsistent with the idea that border ownership assignment represents a surface signal rather than a property tied to a border (Grossberg, 2016; Nakayama et al., 1995; Peterson and Skow, 2008).

### Clustering of border ownership selectivity

Functional clustering is a recurring theme in primate cortical visual areas (Hubel and Livingstone, 1987), and has been reported for several modalities in V4 including orientation, hue, direction of motion, spatial frequency and recently curvature (Hu et al., 2020; Li et al., 2013; Liu et al., 2020; Lu et al., 2018; Roe et al., 2012; Tang et al., 2020; Tanigawa et al., 2010). We find here that border ownership preference is preserved across layers within a vertical penetration, indicating that this modality is also organized in a columnar fashion. Importantly though, border ownership is of a fundamentally different nature than these other modalities: the border ownership of an edge is computed based on stimulus features falling outside the cRF. Combined with our finding that early border ownership signals are computed by deep layer neurons rather than being passed on from upstream areas, this columnar organization has important implications for the functional anatomy. It indicates that there is a systematic asymmetric arrangement of the extra-classical contextual information, which, as argued above, may be provided through horizontal fibers and cortico-cortical feedback. For example, consider a cluster in which neurons prefer that a vertical edge belongs to an object on the left (left column in Figure 7, symbol below left column indicates preferred border ownership). The neurons in this column thus need to receive asymmetric contextual synaptic information favoring the presence of an object to the left of the edge (symbol above the left column in Figure 7). Neurons in another cluster (right column in Figure 7) will instead prefer that the same edge is part of an object on the right, and these neurons will thus necessarily receive asymmetric contextual information of the opposite polarity (symbol above right column). The clustered architecture of preferred border ownership therefore requires clusters of asymmetric contextual information in cortex, such that opposite polarities of contextual information occur in distinct and complementary clusters. The present data indicate that border ownership selectivity initially does not arrive through the feedforward pathway. They thus suggest that the substrate of these clusters consists of clustered asymmetries in the retinotopic information carried by afferents from horizontal fibers and cortical feedback.

## Materials and methods

### Animals

We obtained recordings in two male rhesus macaques (*Macaca mulatta*), age 13 y (animal Z) and age 15 y (animal D). Both animals had no prior experimental history and were housed in separate cages in a primate room with up to six animals of the same species. This study was performed in accordance with the recommendations in the Guide for the Care and Use of Laboratory Animals of the National Institutes of Health. All procedures were approved by the Institutional Animal Care and Use Committee of the Salk Institute for Biological Studies.

### Surgery

Surgical procedures have been described before (Nandy et al., 2017). In brief, in a first surgical session a titanium recording chamber was installed in a craniotomy over the prelunate gyrus, according to stereotactic coordinates derived from anatomical MRI scans from each animal (left hemisphere in animal Z, right hemisphere in animal D). In a second surgical session, the dura mater within the chamber was removed, and replaced with a silicone-based optically clear artificial dura (AD), establishing an optical window over dorsal area V4.

### Electrophysiology

At the beginning of a recording session, a sterile insert consisting of a metal ring covered with a plastic membrane on the bottom was lowered in the chamber. The membrane was perforated to allow insertion of probes. The function of the insert is to stabilize the recording site from cardiopulmonary pulsations. A linear multielectrode probe (32-channel single-shaft acute probes, 100 µm electrode pitch (ATLAS Neuroengineering (Leuven, Belgium)) was mounted on the chamber using a hydraulic microdrive on an adjustable x-y-stage (MO-972A, Narashige (Japan)). The probe was then lowered through the AD over the prelunate gyrus, positioned orthogonally relative to the cortical surface under visual guidance (Zeiss microscope). While monitoring the voltage signals from the electrodes for multiunit activity, the probe was lowered to penetrate the cortical surface. The probe was advanced until multiunit activity was visible on the deepest ∼2600 µm of the probe. Then, the probe was retracted typically by several 100 µm to ease dimpling of the cortex. Between recording sessions, probe position was varied (receptive field eccentricity median 4.87 degrees of visual angle (dva), interquartile range 2.51 dva). Neural signals were recorded extracellularly, filtered and saved using Intan hardware (RHD2132 amplifier chip and RHD2000 amplifier evaluation system, Intan Technologies LLC (Los Angeles, USA)) controlled by a Windows computer.

### Stimulus presentation

Visual stimuli were presented using a LED projector, back-projected on a rear-projection screen that was positioned at a distance of 52 cm from the animal’s eyes (PROPixx, VPixx Technologies (Saint-Bruno, Canada))). The MonkeyLogic software package developed in MATLAB (https://www.brown.edu/Research/monkeylogic/; https://monkeylogic.nimh.nih.gov/) was used for stimulus presentation, behavioral control and recording of eye position. A photodiode was used to measure stimulus timing. Eye position was continuously monitored with an infrared eye tracking system (ISCAN model ETL-200 (Woburn, MA)) and eye traces were saved using MonkeyLogic. Trials were aborted if eye position deviated from the fixation point (threshold typically 1 dva radius).

### Receptive field mapping stimuli

At the beginning of each recording session, receptive field (RF) mapping data was obtained using a subspace reverse correlation approach (Nandy et al., 2017). Stimuli consisted of static modified Gabors, constructed using square-wave instead of sinusoidal gratings, and dark grey rings (80% luminance contrast, diameter 2 dva, thickness 0.25 dva). Grating spatial frequency and phase were such that a single contrast edge was visible centrally in the Gaussian window (grating parameters: 6 orientations, 2 contrast polarities, typically 80% luminance contrast, one half was one of seven colors or greyscale, the other half was always greyscale; window: FWHM 2 dva). The stimuli were presented every 50 or 60 ms while the animal maintained fixation. Each stimulus appeared at a random location selected from a grid sized 25 dva x 25 dva with 1 dva spacing centered at coordinates [7.5 dva; 7.5 dva] in the appropriate visual quadrant. During the recording session, high-gamma filtered voltage waveforms in response to these stimuli were analyzed to estimate the retinotopy of the probe position, and choose location and size for the border ownership stimuli. Detailed receptive fields were calculated for each unit offline after spike sorting and used to verify the proper position of the border ownership stimuli.

### Layer assignment

A CSD mapping procedure on evoked LFP was used to estimate the laminar position of recorded channels (Nandy et al., 2017). Briefly, animals maintained fixation while dark gray ring stimuli were flashed (32 ms stimulus duration, 94% luminance contrast, sized and positioned to fall within the cRF of the probe position). The CSD was calculated as the second spatial derivative of the stimulus-triggered LFP (filtered between 3.3 Hz and 88 Hz) and visualized as spatial maps after smoothing using bicubic 2-D interpolation (Figure 1-figure supplement 1; MATLAB function *interp2* with option *cubic*, the spatial dimension was interpolated at a resolution of 10 µm). Red regions depict current sinks, blue regions depict current sources. As described in more detail in Results, we observed a consistent pattern between different penetrations, strikingly similar to the current sink-source maps reported by other laboratories in behaving macaques beyond V1 (V4: Pettine et al., 2019; V4: Lu et al., 2018; area 36: Takeuchi et al., 2011). Through histological verification Takeuchi et al. (2011) found that the prominent current sink with the shortest latency corresponded to the position of the granular layer. We therefore identified this current sink (current sink indicated by white star, between dashed and dotted white lines in Figure 1F,G; Figure 1-figure supplements 1 and 2) as the granular layer. For each unit, the positions of the electrode contacts (using the five contacts surrounding the one with the largest spike waveform) were weighed by the average peak-to-trough amplitude of the unit’s spike waveform on these contacts, to assign the unit to the depth where its spike waveform is largest. By comparing this position with the range of contacts in the granular layer, we could locate units to superficial, granular or deep layers. Seven penetrations where the CSD map could not be interpreted were excluded from the laminar analyses, and we also required that all units included in the laminar analyses were located ≤2 mm from the most superficial channel with multiunit activity, to minimize the risk of including white matter activity.

### Orientation tuning stimuli

A data set for orientation tuning was obtained using luminance contrast edges similar to those used for RF mapping, but with a circular window, sized and positioned such that the stimulus covered and was centered on the estimated aggregate cRF of the probe (12 orientations; 2 contrast polarities; 54% luminance contrast; 200 ms stimulus duration).

### Border ownership stimuli and task

From the online analysis on the RF mapping data, position, size and color of the border ownership stimulus set were chosen. An isoluminant square was positioned on an isoluminant background, as in prior studies on border ownership (Zhou et al., 2000). There were four basic conditions (Figures 1A,C), consisting of a factorial combination of two square positions and two contrast polarities. Note that this results in two pairs of stimuli, where both scenes in a pair have identical stimulus information inside the cRF (dotted black circles in Figures 1A,C). The two luminance areas in each scene were either both greys or a combination of grey and a color, and luminance contrast was 54%, as in prior studies (Zhou et al., 2000). Square sizes were between 12×12 dva and 18×18 dva. At the beginning of each trial, a small light grey fixation point (0.2 x 0.2 dva, 80% luminance contrast) was presented on a blank grey screen (with luminance set at the geometric mean of the luminances in the border ownership stimulus). After the animal maintained fixation for 400 ms, the border ownership stimulus appeared for 500 ms. Then, for a separate project, the stimulus was then replaced with a stimulus in which the central edge was prolonged to cover the entire screen (with identical colors and luminances) and the other parts of the square removed, for another 1000 ms. Spikes elicited during that time window were not included in the analyses in this paper. The animal received a juice reward if it maintained fixation throughout the trial. Depending on recording time, data was obtained for different orientations or positions. Conditions were played pseudorandomly in counterbalanced blocks such that each condition was played once before repeating conditions. Typically 8-10 repetitions were obtained per condition. Often these stimuli were played at a few different orientations and/or positions, in order to increase the likelihood of proper stimulus placement for most units recorded on the probe (because the precise cRF for each unit was only available offline, after spike sorting). On some days the trials analyzed here were randomly interdigitated with similar trials using stimuli defined for other projects. Some of these trials – not analyzed here – included a condition in which the animal had to saccade to a new fixation position if the fixation point moved.

### Analysis

Data was analyzed in MATLAB (MathWorks, Natick, MA). Circular statistics were computed using the MATLAB toolbox CircStat (Berens, 2009). Statistical tests for the different analyses are described in detail below. Statistical significance was defined as p<0.05.

### Spike sorting

The data was sorted offline using SpyKING CIRCUS (Yger et al., 2018). The clusters resulting from the automatic sorting step were curated manually using the MATLAB GUI provided by the SpyKING CIRCUS software. Well-isolated units were identified based on a well-defined refractory period in the interspike-interval histogram. Multiunit clusters included in the analysis had to pass a criterion for the signal-to-noise ratio: peak-to-peak amplitude of the average waveform had to exceed five times the standard deviation of the signal 5 ms prior to the peak (similar to Kashkoush et al., 2019), after high-pass filtering the data (1-pole butterworth filter, cut-off 300 Hz, implemented using functions *butter* and *filtfilt* in MATLAB).

### Receptive field mapping

To determine the cRF, spikes were counted in a window [30 100] ms after each stimulus onset. The resulting mean counts per stimulus position were transformed to *z* scores by first subtracting the mean of and then dividing by the standard deviation of spike counts occurring in a window of the same size preceding that stimulus position (Keliris et al., 2019). Using the stimulus positions *z* scores were transformed to a spatial map, which was smoothed with a Gaussian filter (MATLAB function *imgaussfilt* with *σ* = 1). The outline of the classical receptive field (cRF) was defined as the contour at *z* = 3 on this smoothed map (calculated using MATLAB function *contourc*).

### Border ownership selectivity

Border ownership responses were obtained by recording evoked responses to stimuli as in Figures 1A,C (see *Border ownership stimuli* above). A unit’s response was evaluated for border ownership selectivity if it passed the following inclusion criteria: 1. average spike rate was ≥1 spike/s for at least one of the four conditions; 2. the evoked spike count for at least one of the four conditions was significantly different from that recorded prior to all trials across conditions (two-sided Wilcoxon rank sum test (MATLAB function *ranksum*) with Bonferroni correction); 3. at least six trials per condition were available; 4. the central edge of the squares in the border ownership stimulus intersected the cRF; 5. the distance between any part of the cRF contour and any part of the non-central edges of the squares in the border ownership stimulus was ≥ 1 dva. Data sets obeying these inclusion criteria were candidate data sets for border ownership selectivity. The border ownership index (*BOI*; Zhou et al., 2000) was then calculated for these data sets, which is defined as

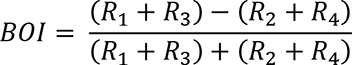

where *R*_*i*_ represents the average spike rate in the window [50 500] ms after stimulus onset for condition *i* (numbering as in Figures 1A,C). Statistical significance of border ownership selectivity was evaluated using a permutation test: a null distribution was created by shuffling the border ownership stimulus labels, separately for each luminance contrast pair, 10000 shuffles), and the p value estimated as the fraction of |*BOI*_shuffled_| that was at least as large as |*BOI*|. A data set was defined to be border ownership selective if p<0.05 (with Bonferroni correction if multiple orientations or positions were available for the same unit), and a unit was defined to be border ownership selective if it had at least one border ownership selective data set.

### Time course of evoked response

For the time course of evoked activity of border ownership selective units (Figures 2A-D), each unit contributed one data set, i.e. the border ownership selective data set – as defined above – for which |*BOI*| was maximal. For each of these, the time courses of the responses to the preferred side of border ownership (side resulting in the highest spike rate) and to the non-preferred side of border ownership (side resulting in the lowest spike rate) were calculated separately (respectively solid red lines and dashed blue lines Figures 2A-D) as follows. Spike trains were rounded to 0.1 ms resolution and convolved with a postsynaptic kernel *K(t)* (Thompson et al., 1996)

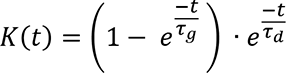

where *τ*_*g*_ = 1 ms and *τ*_*d*_ = 20 ms. The resulting traces were averaged per condition, and then across both contrast polarities. These average traces were normalized for each unit’s evoked response by dividing them by the average value across conditions in the window [50 500] ms after stimulus onset. The mean ± s.e.m. of the resulting functions across units are shown in Figures 2A-D. The latency of statistical difference between the functions to the preferred and the non-preferred functions (asterisks in Figure 2A-D; Figure 2-figure supplement 1) was defined as the first time after which the functions differed statistically (Wilcoxon sign rank test p<0.05) for 20 consecutive milliseconds.

Border ownership index functions (Figure 2E) were defined as the difference between the response function for the preferred side of border ownership and the function for the non-preferred side of border ownership, divided by their sum. Latency of these functions was defined as the earliest crossing of a fixed threshold that was followed by values above the threshold for 20 consecutive milliseconds. The threshold used was derived from shuffled data, by shuffling the stimulus labels for each laminar compartment (1000 shuffles) and finding the lowest value for which <1% of shuffles resulted in a defined latency. Since these values depend on sample size, the highest value across compartments was used as threshold, so that the different functions could be timed using the same threshold. Confidence intervals (95%) were calculated on these latencies using a bootstrap approach with the bias corrected and accelerated percentile method (MATLAB function *bootci*, 2000 bootstraps). The latencies were statistically compared between laminar compartments using a bootstrap approach similar to other studies (Self et al., 2019), by computing the difference in latency between bootstrap samples of different compartments and estimating p as the fraction of samples on which the difference was less than or equal to zero (one-sided test).

Spike response functions to small rings in the cRF (Figure 3) were computed similarly as the response functions to border ownership stimuli (Figure 2A-D). Responses were normalized by dividing them by the peak response for each unit. Latency was defined using a threshold halfway between baseline and peak (0.435), and confidence intervals and statistical tests were computed in the same way as for the border ownership index functions.

### Border ownership reliability and latency

We also evaluated the time course of border ownership selectivity using the reliability metric (Zhou et al., 2000), which reflects the trial-to-trial reliability of encoding border ownership. This metric was computed for the border ownership selective units in each laminar compartment, in 100 ms sliding windows (1 ms steps). For each unit, 10000 sets of four spike trains were generated, where each set contained one random spike train from each of the four conditions in Figures 1A,C. For each window position, spikes were counted for each spike train in each set. Border ownership reliability (*BOR*) for a unit at a particular window position was defined as

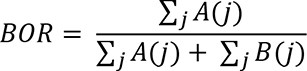

where *j* corresponds to the index of all spike train sets for the unit. *A*(*j*) and *B*(*j*) indicate whether the sign of the spike count difference between border ownership conditions for spike train set *j* is respectively the same or opposite compared to the unit’s preferred side of border ownership:

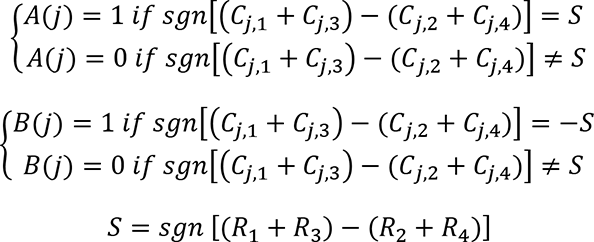

where *C*_*j*,*i*_ represents the window spike count for condition *i* in spike train set *j*, *R*_*i*_ is the average spike rate for condition *i* (for the interval [50 500] ms after stimulus onset), and *sgn* is the sign function. For each window position for each unit, *BOR* was only computed if there were at least 10 spikes across conditions. For every window position the mean was calculated across units per laminar compartment, resulting in the *BOR* functions shown in Figure 2F. The abscissa in Figure 2F corresponds to the position of the right edge of the window, i.e. it indicates the latest spike times that may have determined *BOR* for that window position (and thus represents a conservative estimate for the latency). Definition of latency, computation of confidence intervals and statistical tests used to compare latencies between layers were similar to those for the border ownership index functions.

### Clustering of preferred side of border ownership

For a given penetration, for each orientation, the border ownership selective data set with the highest |*BOI*| was selected for each unit. Then from this group of data sets the largest subgroup that shared the same edge orientation and position were retained for analysis. Each unit could thus maximally contribute one data set. The preferred side of border ownership was then determined for all these data sets (example penetrations are shown in Figure 4C). The proportion of units sharing the most common preferred side was calculated for each penetration. The average of these proportions across penetrations (*P_pref_*) was compared with a null distribution generated by randomly assigning the preferred side of border ownership for each data set (2000 randomizations; i.e. a binomial process with chance of success = 0.5). The p value was estimated as the fraction of the null distribution for which *P_pref_* was at least as large as the actual data.

### Orientation tuning

Orientation selectivity was evaluated using a Kruskal-Wallis test (MATLAB function *kruskalwallis*) on the evoked spike counts in the analysis window (between 30 ms and 200 after stimulus onset) from the orientation tuning data set, using orientation as the grouping variable (Pettine et al., 2019), and defined as p<0.05. All units for which the center of the contrast edge in the orientation data set was positioned in the cRF, and the z-scored spike rate for the orientation with the highest rate was ≥ 3 were included. *Z*-scores were calculated by first subtracting the mean of and then dividing by the standard deviation of the spike rate in a window preceding the first stimulus in the trial, equal in duration to the analysis window. For each orientation-selective unit *i*, the response to each orientation *j* is summarized as 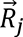, which has a direction 2*j* (because orientation has a period of 180°) and magnitude equal to the spike rate. The resultant 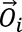 (vectors shown as solid lines in Figure 5) for all 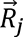 is then calculated. The magnitude of orientation-selectivity and the preferred orientation of unit *i* were defined respectively as the magnitude and as the direction divided by 2 (because of the definition of direction of 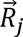) of 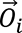.

For each penetration, the aggregate preferred orientation was defined as the direction divided by 2 of the resultant vector 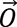 of all 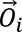 on that penetration (direction of 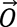 is indicated by blue dashed lines in Figure 5). Statistical significance of the aggregate preferred orientation was assessed by randomizing the directions of 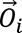 (2000 shuffles), calculating the null distribution 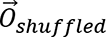, and estimating the p-value as the fraction of 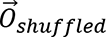 for which the magnitude was at least as large as that of 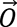.

The independence between the fractions of respectively orientation-selective and border ownership-selective units (Table) was assessed using a Chi square test (MATLAB function *crosstab*).

For units that were border ownership selective to multiple orientations, the circular mean of border ownership selective orientations (ordinate in Figure 6C) was calculated as the direction divided by 2 of the resultant vector of all vectors 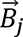 that have a direction 2*j* and a magnitude equal to |*BOI*|, for all orientations *j*. The shortest distance of the data points in Figure 6C to the identity line (Figure 6D) was analyzed by comparing the mean to a null distribution generated by shuffling the values for the preferred edge orientation and the circular mean of border ownership-selective orientations, and calculating the mean of the shortest distance to the identity line (2000 shuffles). The p value was estimated as the fraction of the null distribution that was as small or smaller than the observed value.

## Acknowledgements

This work was supported by a Fellowship from the George E. Hewitt Foundation for Medical Research (to T.P.F.), a NARSAD Young Investigator Grant from the Brain & Behavior Research Foundation (to T.P.F.), NIH Grant K99EY031795 (to T.P.F.), NIH Grant P30-EY0190005 (Salk Core Grant for Vision Research) and the Fiona and Sanjay Jha Chair in Neuroscience (to J.H.R.). We thank Dr. Edward Callaway, Dr. Anirvan Nandy and Dr. Zachary Davis for helpful discussions. We thank Dr. Mathias LeBlanc, Dr. Sean Adams and Ms. Catherine Williams for excellent animal care.

---

**Figure 1 – figure supplement 1.**
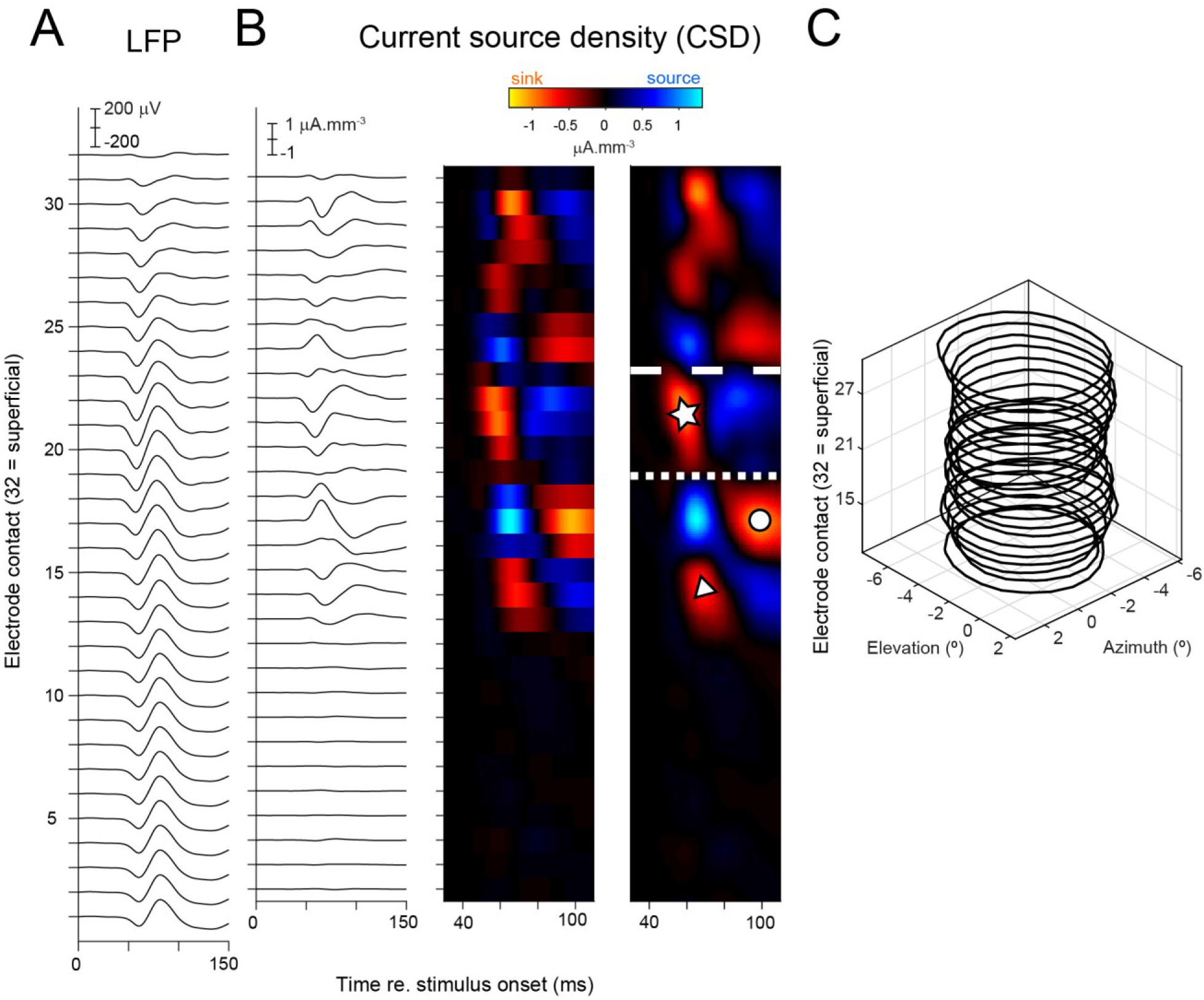
Construction of current source density (CSD) map. Data shown from an example penetration. (**A**) Average local field potential from each electrode contact in response to small rings positioned in the cRF. (**B**) CSD is computed as the second spatial derivative of the LFP. Left panel: CSD traces. Negative values correspond to current sinks, positive values to current sources. Middle panel: CSD traces plotted on a color scale. Right panel: smoothed CSD map (Methods). Symbols and lines between compartments drawn as in Figures 1F,1G. (**C**) cRF contours for different electrodes from this penetration (contours drawn at *z* = 3).

**Figure 1 – figure supplement 2.**
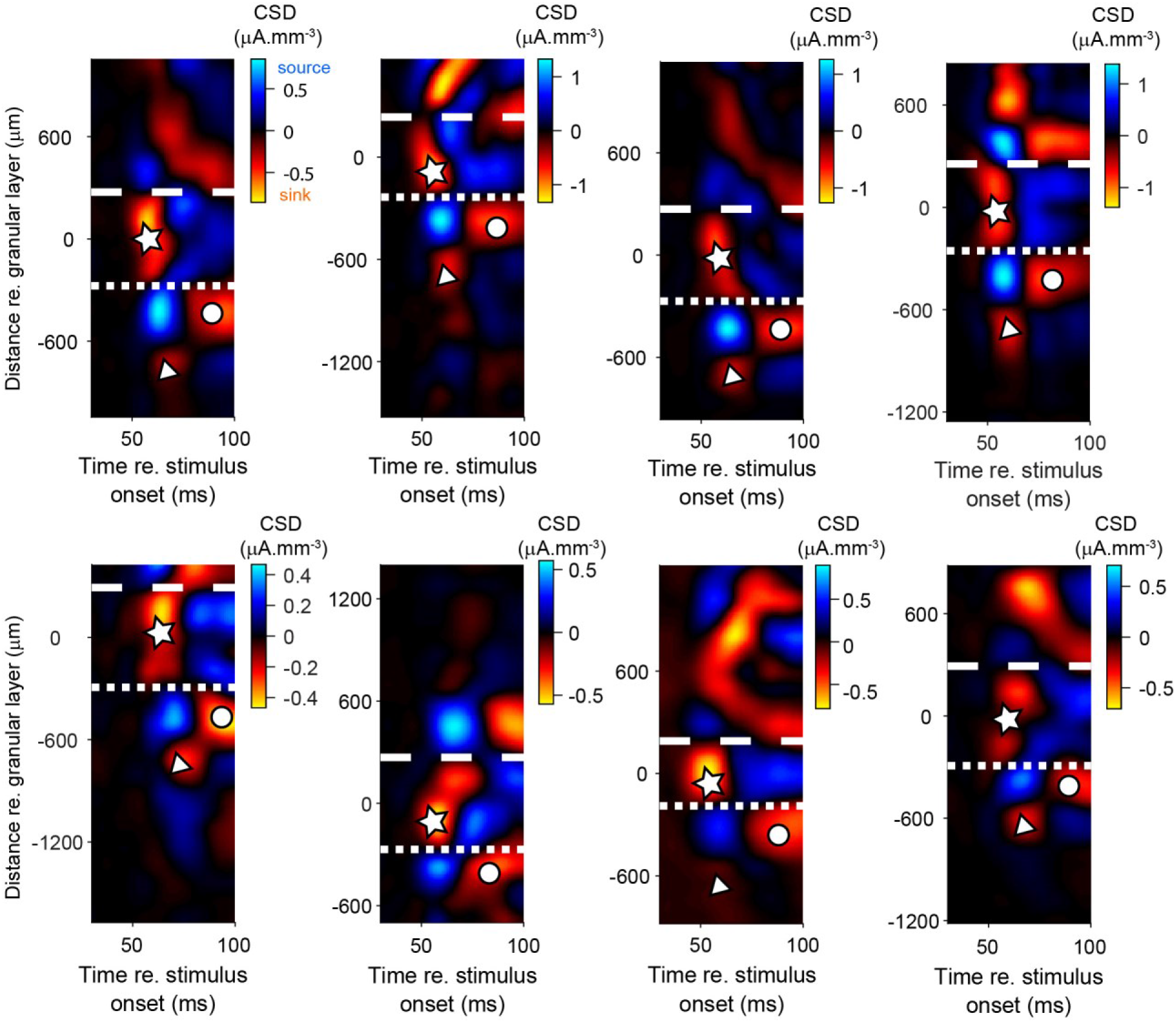
Additional examples of current source density (CSD) maps. Each panel shows a CSD map from a different penetration. Reddish colors are current sinks, blueish colors are current sources. Symbols as in Figures 1F,1G. Sink-source patterns are strikingly similar to those reported in the behaving macaque in V4 by other laboratories (Pettine et al., 2019, their Figure 1C; Lu et al., 2018, their Figure S4B), and in another downstream cortical area (area 36: Takeuchi et al., 2011, their Figure S1). Top row panels are from penetrations from animal D, bottom panels are from penetrations from animal Z.

**Figure 2 – figure supplement 1.**
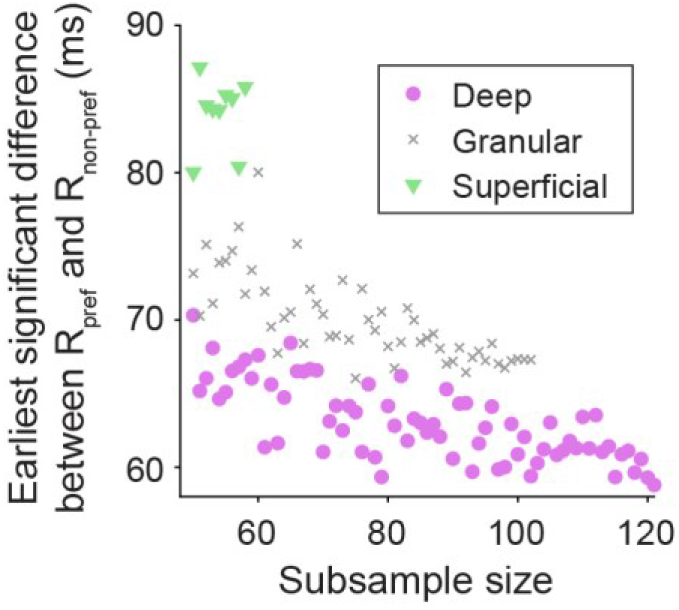
Differences in sample size do not explain differences in latency between layers. The latency of significant difference between responses to preferred and non-preferred sides of border ownership was determined as in Figure 2B-D, for different subsample sizes (subsampling well-isolated units without replacement). Each subsample size was sampled five times and the average latency is plotted. The result shows that while the total number of units in each layer did differ, the shorter latency for deep layer units than for granular or superficial layer units consistently appeared at all subsample sizes (consistent with data shown in Figure 2B-D), and is not explained by differences in sample size between compartments.

**Figure 2 – figure supplement 2.**
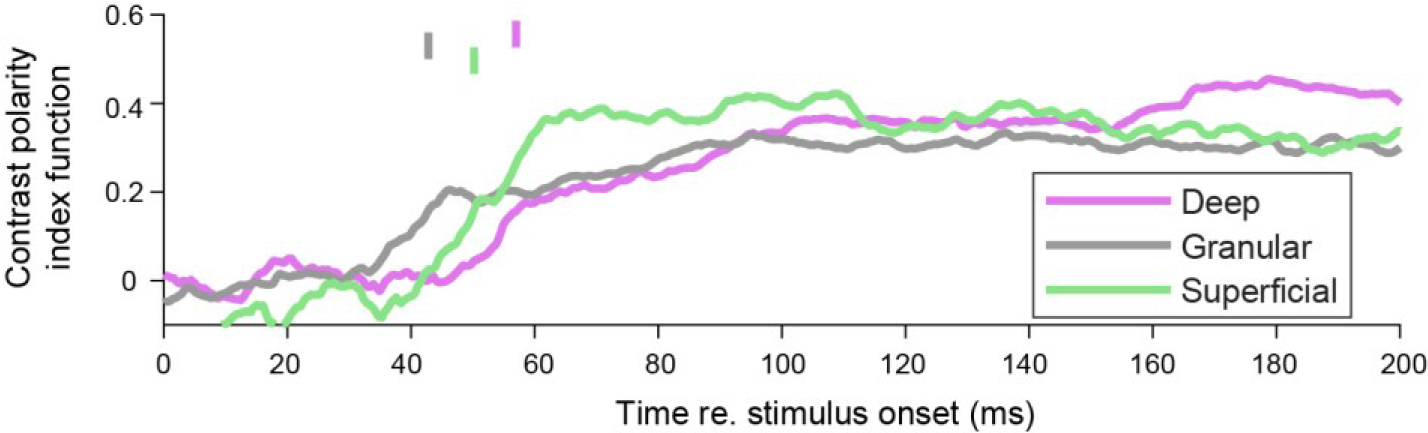
Selectivity for contrast polarity does not occur earliest in the deep layers. Same analysis as Figure 2E, but for contrast polarity instead of border ownership, for the same units as in Figure 2E. Latency at same threshold as in Figure 2E: deep: 57.0 ms, 95% CI [49.9 75.0]; granular: 42.8 ms, 95% CI [35.4 60.8]; superficial: 50.2, 95%CI [40.4 56.4]. Granular vs. deep: bootstrap procedure p=0.14. Granular vs. superficial: p=0.12. That border ownership units are often selective for contrast polarity is consistent with prior work (Zhou et al., 2000).

**Figure 4–figure supplement 1.**
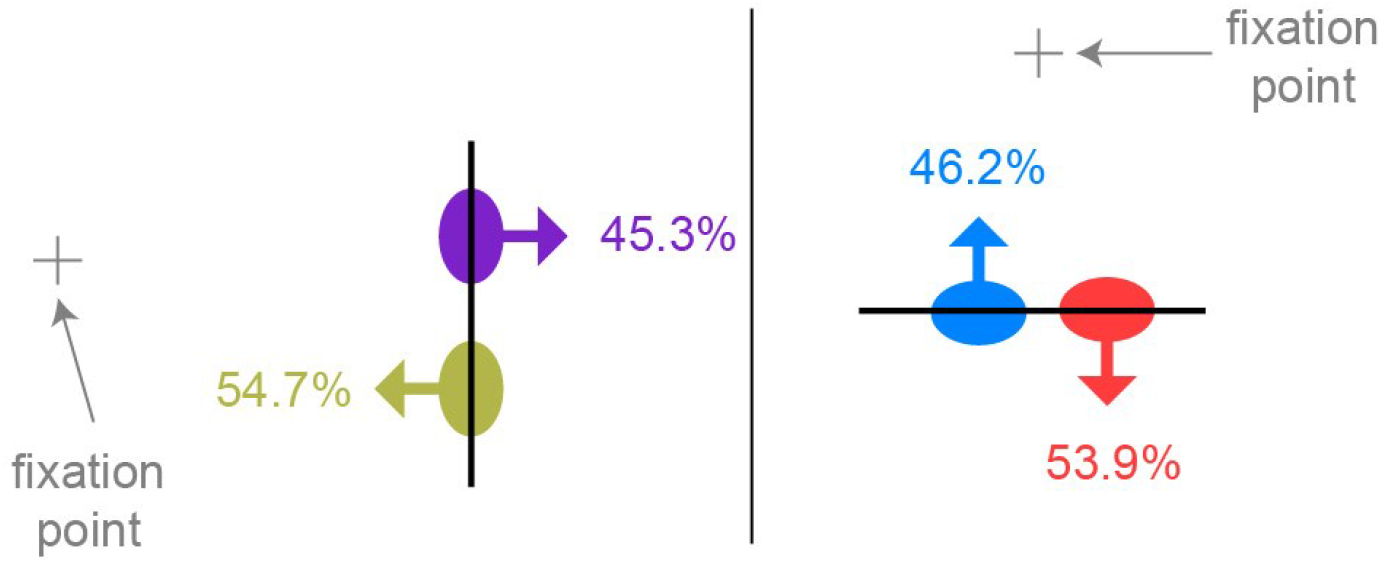
Similar to Figure 4A, but expressing preferred side of border ownership relative to the position of the fixation point. Together with Figure 4A, this indicates that similarly sized populations of border ownership units with opposite preferences occur in the same visual quadrant. Vertical edge: n = 64 well-isolated units selective for border ownership; horizontal edge: n = 78 well-isolated units selective for border ownership.

**Figure 6–figure supplement 1.**
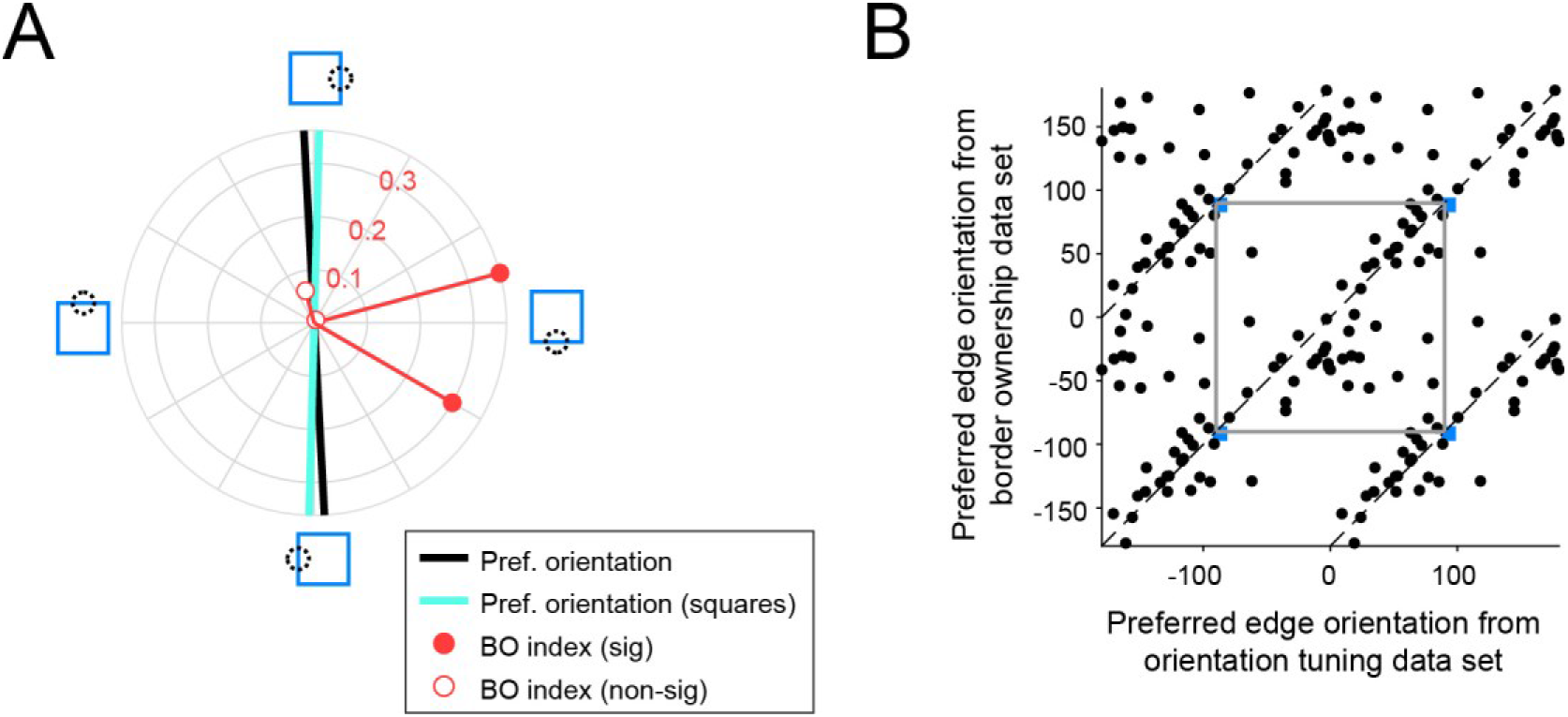
Consistent orientation tuning for isolated edges and for edges that are part of squares. (**A**) Same plot as in Figure 6A (middle), but including preferred edge orientation derived from the border ownership stimulus set (cyan line). (**B**) Preferred edge orientation derived from the border ownership stimulus set (edges that are part of squares) plotted against preferred edge orientation derived from the orientation tuning data set (isolated contrast edges). As in Figure 6C, data points are plotted repeatedly every 180° for these periodic variables and two periods are shown. The grey square outlines an area corresponding to one period for both variables. Dashed lines indicate identical values. Blue square indicates the unit shown in A. Values are significantly clustered along the identity line (randomization test p=0.0005; n = 51 units).

## References

Angelucci, A., Bijanzadeh, M., Nurminen, L., Federer, F., Merlin, S., Bressloff, P.C., 2017. Circuits and Mechanisms for Surround Modulation in Visual Cortex. Annu. Rev. Neurosci. 40, 425–451. https://doi.org/10.1146/annurev-neuro-072116-031418

Berens, P., 2009. CircStat: A MATLAB toolbox for circular statistics. J. Stat. Softw. 31, 1–21. https://doi.org/10.18637/jss.v031.i10

Binzegger, T., Douglas, R.J., Martin, K.A.C., 2004. A quantitative map of the circuit of cat primary visual cortex. J. Neurosci. 24, 8441–8453. https://doi.org/10.1523/JNEUROSCI.1400-04.2004

Briggs, F., Callaway, E.M., 2005. Laminar patterns of local excitatory input to layer 5 neurons in macaque primary visual cortex. Cereb. Cortex N. Y. N 1991 15, 479–488. https://doi.org/10.1093/cercor/bhh154

Callaway, E.M., 1998. Prenatal development of layer-specific local circuits in primary visual cortex of the macaque monkey. J. Neurosci. 18, 1505–1527.

Craft, E., Schütze, H., Niebur, E., von der Heydt, R., 2007. A neural model of figure-ground organization. J. Neurophysiol. 97, 4310–4326. https://doi.org/10.1152/jn.00203.2007

Douglas, R.J., Martin, K.A., 2004. Neuronal circuits of the neocortex. Annu. Rev. Neurosci. 27, 419–451. https://doi.org/10.1146/annurev.neuro.27.070203.144152

Douglas, R.J., Martin, K.A., 2007. Mapping the Matrix: The Ways of Neocortex. Neuron 56,226–238. https://doi.org/10.1016/j.neuron.2007.10.017

Fries, W., 1984. Cortical projections to the superior colliculus in the macaque monkey: a retrograde study using horseradish peroxidase. J. Comp. Neurol. 230, 55–76. https://doi.org/10.1002/cne.902300106

Gattass, R., Sousa, A.P., Mishkin, M., Ungerleider, L.G., 1997. Cortical Projections of Area V2 in the Macaque. Cereb. Cortex 7, 110–129. https://doi.org/10.1093/cercor/7.2.110

Gattass, R., Galkin, T.W., Desimone, R., Ungerleider, L.G., 2014. Subcortical connections of area V4 in the macaque. J. Comp. Neurol. 522, 1941–1965. https://doi.org/10.1002/cne.23513

Ghose, G.M., Ts’o, D.Y., 1997. Form processing modules in primate area V4. J. Neurophysiol. 77, 2191–2196. https://doi.org/10.1152/jn.1997.77.4.2191

Girard, P., Hupé, J.M., Bullier, J., 2001. Feedforward and feedback connections between areas V1 and V2 of the monkey have similar rapid conduction velocities. J. Neurophysiol. 85, 1328–1331. https://doi.org/10.1152/jn.2001.85.3.1328

Grossberg, S., 2016. Cortical dynamics of figure-ground separation in response to 2D pictures and 3D scenes: how V2 combines border ownership, stereoscopic cues, and Gestalt grouping rules. Front. Psychol. 6. https://doi.org/10.3389/fpsyg.2015.02054

Harris, K.D., Shepherd, G.M., 2015. The neocortical circuit: themes and variations. Nat. Neurosci. 18, 170–181. https://doi.org/10.1038/nn.3917

Hesse, J.K., Tsao, D.Y., 2016. Consistency of border-ownership cells across artificial stimuli, natural stimuli, and stimuli with ambiguous contours. J. Neurosci. 36, 11338–11349. https://doi.org/10.1523/JNEUROSCI.1857-16.2016

Hu, B., Niebur, E., 2017. A recurrent neural model for proto-object based contour integration and figure-ground segregation. J. Comput. Neurosci. 43, 227–242. https://doi.org/10.1007/s10827-017-0659-3

Hu, J.M., Song, X.M., Wang, Q., Roe, A.W., 2020. Curvature domains in V4 of macaque monkey. eLife 9. https://doi.org/10.7554/eLife.57261

Hubel, D. H., Livingstone, M. S. (1987). Segregation of form, color, and stereopsis in primate area 18. J. Neurosci. 7, 3378–3415.

Hubel, D.H., Wiesel, T.N., Stryker, M.P., 1978. Anatomical demonstration of orientation columns in macaque monkey. J. Comp. Neurol. 177, 361–379. https://doi.org/10.1002/cne.901770302

Jiang, R., Andolina, I.M., Li, M., Tang, S., 2021. Clustered functional domains for curves and corners in cortical area V4. Elife 10:e63798. doi: https://doi.org/10.7554/eLife.63798

Kashkoush, A.I., Gaunt, R.A., Fisher, L.E., Bruns, T.M., Weber, D.J., 2019. Recording single- and multi-unit neuronal action potentials from the surface of the dorsal root ganglion. Sci. Rep. 9, 2786. https://doi.org/10.1038/s41598-019-38924-w

Keliris, G.A., Li, Q., Papanikolaou, A., Logothetis, N.K., Smirnakis, S.M., 2019. Estimating average single-neuron visual receptive field sizes by fMRI. Proc. Natl. Acad. Sci. U. S. A. 116, 6425–6434. https://doi.org/10.1073/pnas.1809612116

Koffka, K., 1935. Principles of Gestalt psychology, Principles of Gestalt psychology. Harcourt, Brace, Oxford, England.

Li, P., Zhu, S., Chen, M., Han, C., Xu, H., Hu, J., Fang, Y., Lu, H.D., 2013. A motion direction preference map in monkey V4. Neuron 78, 376–388. https://doi.org/10.1016/j.neuron.2013.02.024

Liu, Y., Li, M., Zhang, X., Lu, Y., Gong, H., Yin, J., Chen, Z., Qian, L., Yang, Y., Andolina, I.M., Shipp, S., Mcloughlin, N., Tang, S., Wang, W., 2020. Hierarchical representation for chromatic processing across macaque V1, V2, and V4. Neuron 108, 538-550.e5. https://doi.org/10.1016/j.neuron.2020.07.037

Lübke, J., Egger, V., Sakmann, B., Feldmeyer, D., 2000. Columnar Organization of Dendrites and Axons of Single and Synaptically Coupled Excitatory Spiny Neurons in Layer 4 of the Rat Barrel Cortex. J Neurosci 20, 5300–5311. https://doi.org/10.1523/JNEUROSCI.20-14-05300.2000

Lu, Y., Yin, J., Chen, Z., Gong, H., Liu, Y., Qian, L., Li, X., Liu, R., Andolina, I.M., Wang, W., 2018. Revealing detail along the visual hierarchy: neural clustering preserves acuity from V1 to V4. Neuron 98, 417–428.e3. https://doi.org/10.1016/j.neuron.2018.03.009

Lund, J.S., Boothe, R.G., 1975. Interlaminar connections and pyramidal neuron organisation in the visual cortex, area 17, of the Macaque monkey. J. Comp. Neurol. 159, 305–334. https://doi.org/10.1002/cne.901590303

Lund, J.S., Yoshioka, T., Levitt, J.B., 1993. Comparison of intrinsic connectivity in different areas of macaque monkey cerebral cortex. Cereb. Cortex N. Y. N 1991 3, 148–162. https://doi.org/10.1093/cercor/3.2.148

Markov, N.T., Misery, P., Falchier, A., Lamy, C., Vezoli, J., Quilodran, R., Gariel, M.A., Giroud, P., Ercsey-Ravasz, M., Pilaz, L.J., Huissoud, C., Barone, P., Dehay, C., Toroczkai, Z., Van Essen, D.C., Kennedy, H., Knoblauch, K., 2011. Weight consistency specifies regularities of macaque cortical networks. Cereb. Cortex N. Y. N 1991 21, 1254–1272. https://doi.org/10.1093/cercor/bhq201

Markov, N.T., Vezoli, J., Chameau, P., Falchier, A., Quilodran, R., Huissoud, C., Lamy, C., Misery, P., Giroud, P., Ullman, S., Barone, P., Dehay, C., Knoblauch, K., Kennedy, H., 2014. Anatomy of hierarchy: feedforward and feedback pathways in macaque visual cortex. J. Comp. Neurol. 522, 225–259. https://doi.org/10.1002/cne.23458

Maunsell, J.H., van Essen, D.C., 1983. The connections of the middle temporal visual area (MT) and their relationship to a cortical hierarchy in the macaque monkey. J. Neurosci. 3, 2563–2586. https://doi.org/10.1523/JNEUROSCI.03-12-02563.1983

Mitzdorf, U., 1985. Current source-density method and application in cat cerebral cortex: investigation of evoked potentials and EEG phenomena. Physiol. Rev. 65, 37–100. https://doi.org/10.1152/physrev.1985.65.1.37

Nakayama, K., He, Z.J., Shimojo, S., 1995. Visual surface representation: a critical link between lower-level and higher-level vision, in: An Invitation to Cognitive Science. Visual Cognition: An Invitation to Cognitive Science. The MIT Press, pp. 1–70.

Nandy, A.S., Nassi, J.J., Reynolds, J.H., 2017. Laminar organization of attentional modulation in macaque visual area V4. Neuron 93, 235–246. https://doi.org/10.1016/j.neuron.2016.11.029

O’Herron, P., von der Heydt, R., 2009. Short-term memory for figure-ground organization in the visual cortex. Neuron 61, 801–809. https://doi.org/10.1016/j.neuron.2009.01.014

Otero-Millan, J., Troncoso, X.G., Macknik, S.L., Serrano-Pedraza, I., Martinez-Conde, S., 2008. Saccades and microsaccades during visual fixation, exploration, and search: Foundations for a common saccadic generator. J. Vis. 8, 21–21. https://doi.org/10.1167/8.14.21

Peterson, M.A., Skow, E., 2008. Inhibitory competition between shape properties in figure-ground perception. J. Exp. Psychol. Hum. Percept. Perform. 34, 251–67. https://doi.org/10.1037/0096-1523.34.2.251

Pettine, W.W., Steinmetz, N.A., Moore, T., 2019. Laminar segregation of sensory coding and behavioral readout in macaque V4. Proc. Natl. Acad. Sci. U. S. A. 116, 14749–14754. https://doi.org/10.1073/pnas.1819398116

Raiguel, S.E., Xiao, D.K., Marcar, V.L., Orban, G.A., 1999. Response Latency of Macaque Area MT/V5 Neurons and Its Relationship to Stimulus Parameters. J Neurophys. 82,1944–1956. https://doi.org/10.1152/jn.1999.82.4.1944

Rockland, K.S., Lund, J.S., 1983. Intrinsic laminar lattice connections in primate visual cortex. J. Comp. Neurol. 216, 303–318. https://doi.org/10.1002/cne.902160307

Rockland, K.S., Pandya, D.N., 1979. Laminar origins and terminations of cortical connections of the occipital lobe in the rhesus monkey. Brain Res. 179, 3–20.

Rockland, K.S., Saleem, K.S., Tanaka, K., 1994. Divergent feedback connections from areas V4 and TEO in the macaque. Vis. Neurosci. 11, 579–600. https://doi.org/10.1017/S0952523800002480

Roe, A.W., Chelazzi, L., Connor, C.E., Conway, B.R., Fujita, I., Gallant, J.L., Lu, H., Vanduffel, W., 2012. Toward a unified theory of visual area V4. Neuron 74, 12–29. https://doi.org/10.1016/j.neuron.2012.03.011

Rothkopf, C.A., Ballard, D.H., Hayhoe, M.M., 2007. Task and context determine where you look. J. Vis. 7, 16–16. https://doi.org/10.1167/7.14.16

Rubin, E., 1921. Visuell wahrgenommene Figuren: Studien in psychologischer Analyse. Copenhagen: Gyldendals.

Sakai, K., Nishimura, H., 2006. Surrounding suppression and facilitation in the determination of border ownership. J. Cogn. Neurosci. 18, 562–579. https://doi.org/10.1162/jocn.2006.18.4.562

Sakai, K., Nishimura, H., Shimizu, R., Kondo, K., 2012. Consistent and robust determination of border ownership based on asymmetric surrounding contrast. Neural Netw. 33, 257– 274. https://doi.org/10.1016/j.neunet.2012.05.006

Self, M.W., Jeurissen, D., van Ham, A.F., van Vugt, B., Poort, J., Roelfsema, P.R., 2019. The segmentation of proto-objects in the monkey primary visual cortex. Curr. Biol. CB 29, 1019–1029.e4. https://doi.org/10.1016/j.cub.2019.02.016

Supèr, H., Romeo, A., Keil, M., 2010. Feed-forward segmentation of figure-ground and assignment of border-ownership. PloS One 5, e10705. https://doi.org/10.1371/journal.pone.0010705

Takahashi, N., Oertner, T.G., Hegemann, P., Larkum, M.E., 2016. Active cortical dendrites modulate perception. Science 354, 1587–1590. https://doi.org/10.1126/science.aah6066

Takeuchi, D., Hirabayashi, T., Tamura, K., Miyashita, Y., 2011. Reversal of Interlaminar Signal Between Sensory and Memory Processing in Monkey Temporal Cortex. Science 331, 1443–1447. https://doi.org/10.1126/science.1199967

Tang, R., Song, Q., Li, Y., Zhang, R., Cai, X., Lu, H.D., 2020. Curvature-processing domains in primate V4. eLife 9, e57502. https://doi.org/10.7554/eLife.57502

Tanigawa, H., Lu, H.D., Roe, A.W., 2010. Functional organization for color and orientation in macaque V4. Nat. Neurosci. 13, 1542–1548. https://doi.org/10.1038/nn.2676

Thompson, K.G., Hanes, D.P., Bichot, N.P., Schall, J.D., 1996. Perceptual and motor processing stages identified in the activity of macaque frontal eye field neurons during visual search. J. Neurophysiol. 76, 4040–4055. https://doi.org/10.1152/jn.1996.76.6.4040

Ungerleider, L.G., Desimone, R., 1986. Cortical connections of visual area MT in the macaque. J. Comp. Neurol. 248, 190–222. https://doi.org/10.1002/cne.902480204

Vanduffel, W., Tootell, R.B.H., Schoups, A.A., Orban, G.A., 2002. The organization of orientation selectivity throughout macaque visual cortex. Cereb. Cortex 12, 647–662. https://doi.org/10.1093/cercor/12.6.647

von der Heydt, R., 2015. Figure-ground organization and the emergence of proto-objects in the visual cortex. Front. Psychol. 6, 1695. https://doi.org/10.3389/fpsyg.2015.01695

Wagatsuma, N., Sakai, K., 2017. Modeling the time-course of responses for the border ownership selectivity based on the integration of feedforward signals and visual cortical interactions. Front. Psychol. 7. https://doi.org/10.3389/fpsyg.2016.02084

Xu, X., Olivas, N.D., Ikrar, T., Peng, T., Holmes, T.C., Nie, Q., Shi, Y. (2016). Primary visual cortex shows laminar-specific and balanced circuit organization of excitatory and inhibitory synaptic connectivity. J. Physiol. 1891–1910. https://doi.org/10.1113/JP271891

Yazdanbakhsh, A., Livingstone, M. S. (2006). End stopping in V1 is sensitive to contrast. Nature Neuroscience, 9(5), 697–702. https://doi.org/10.1038/nn1693

Yger, P., Spampinato, G.L., Esposito, E., Lefebvre, B., Deny, S., Gardella, C., Stimberg, M., Jetter, F., Zeck, G., Picaud, S., Duebel, J., Marre, O., 2018. A spike sorting toolbox for up to thousands of electrodes validated with ground truth recordings in vitro and in vivo. eLife 7, e34518. https://doi.org/10.7554/eLife.34518

Yoshioka, T., Levitt, J.B., Lund, J.S., 1992. Intrinsic lattice connections of macaque monkey visual cortical area V4. J. Neurosci. 12, 2785–2802.

Zarrinpar, A., Callaway, E.M., 2016. Functional local input to layer 5 pyramidal neurons in the rat visual cortex. Cereb. Cortex N. Y. N 1991 26, 991–1003. https://doi.org/10.1093/cercor/bhu268

Zhang, N.R., von der Heydt, R., 2010. Analysis of the context integration mechanisms underlying figure–ground organization in the visual cortex. J. Neurosci. 30, 6482–6496. https://doi.org/10.1523/JNEUROSCI.5168-09.2010

Zhaoping, L., 2005. Border ownership from intracortical interactions in visual area v2. Neuron 47, 143–153. https://doi.org/10.1016/j.neuron.2005.04.005

Zhou, H., Friedman, H.S., von der Heydt, R., 2000. Coding of border ownership in monkey visual cortex. J. Neurosci. 20, 6594–6611.

